# Quantifying the fraction of new mutations that are recessive lethal

**DOI:** 10.1101/2022.04.22.489225

**Authors:** Emma E. Wade, Christopher C. Kyriazis, Maria Izabel A. Cavassim, Kirk E. Lohmueller

## Abstract

The presence and impact of recessive lethal mutations has been widely documented in diploid outcrossing species. However, precise estimates in different species of the proportion of mutations that are recessive lethal remain limited. Here, we attempt to quantify the fraction of new mutations that are recessive lethal using Fit*∂*a*∂*i, a commonly-used method for inferring the distribution of fitness effects (DFE) using the site frequency spectrum. Using simulations, we demonstrate that Fit*∂*a*∂*i cannot accurately estimate the fraction of recessive lethal mutations, as expected given that Fit*∂*a*∂*i assumes that all mutations are additive by default. Consistent with the idea that mis-specification of the dominance model can explain this performance, we find that Fit*∂*a*∂*i can accurately infer the fraction of additive lethal mutations. Moreover, we demonstrate that in both additive and recessive cases, inference of the deleterious non-lethal portion of the DFE is minimally impacted by a small proportion (<10%) of lethal mutations. Finally, as an alternative approach to estimate the proportion of mutations that are recessive lethal, we employ models of mutation-selection-drift balance using existing genomic parameters and segregating recessive lethals estimates for humans and *Drosophila melanogaster*. In both species, we find that the segregating recessive lethal load can be explained by a very small fraction (<1%) of new nonsynonymous mutations being recessive lethal. Our results refute recent assertions of a much higher recessive lethal mutation fraction (4-5%), while highlighting the need for additional information on the joint distribution of selection and dominance coefficients.

## Introduction

From the early days of genetics, it was noted that it was often not possible to make a chromosome homozygous in a *Drosophila* cross. This inability to make a chromosome homozygous was taken as evidence that it carried a recessive lethal mutation in the heterozygous state that resulted in death or sterility when made homozygous (Dobzhansky, Spassky, and Spassky 1954; Dubinin 1946). Through further study, approximately 20-40% of *Drosophila melanogaster* autosomes sampled from natural populations could not be made homozygous, implying ∼1.6 recessive lethal mutations per diploid autosome (Simmons and Crow 1977). Similar estimates of ∼1-2 recessive lethals per diploid genome have also been obtained from natural zebrafish and bluefin killifish populations, despite these species having genome sizes an order of magnitude larger than *D. melanogaster* (McCune et al. 2002; Halligan and Keightley 2003). Together, these studies suggest that the recessive lethal load may be relatively constrained across species.

Obtaining estimates of the number of segregating recessive lethals in humans has required more indirect approaches, given that humans are not amenable to experimentation. In the 1950s, Morton et al. found that the rate of juvenile mortality in humans increased with increasing relatedness between an individual’s parents (Morton, Crow, and Muller 1956). Such studies have revealed that humans typically carry 1.4 diploid lethal equivalents that, if made homozygous, would result in a genetic death (Bittles and Neel 1994). This estimate may therefore represent an upper bound on the number of recessive lethals, as the total number of lethal equivalents in a genome represents the cumulative effect of all recessive deleterious mutations in a genome (Morton, Crow, and Muller 1956). More recently, an elegant study combined the observed incidence of recessive lethal disease in a founder population combined with pedigree records and simulations to more directly estimate the number of recessive lethal mutations carried by the founders (Gao et al. 2015). They inferred that each founder carried ∼0.6 recessive lethal mutations, however, they note that this may be a slight underestimate for various reasons (Gao et al. 2015). Thus, available evidence suggests that humans have a recessive lethal load that is roughly consistent with estimates from other species.

Separately from this work, a growing number of studies have attempted to use genetic variation data to estimate the distribution of fitness effects of new mutations (DFE) in humans and other species. This distribution quantifies the expected effects on fitness of new mutations entering a population, including lethal mutations. In other words, it is the distribution of selection coefficients (*s*) for new mutations in a particular type of sites in the genome (e.g., nonsynonymous mutations). These approaches rely on genetic variation data from resequencing of individuals from natural populations, which is summarized by the site frequency spectrum (SFS), i.e., the numbers of variants with particular frequencies in the sample. The parameters of the DFE are then estimated using population genetic models of mutation, selection, and demography that match this observed SFS. This approach has been applied to infer the DFE from numerous taxa including prokaryotes (Cavassim et al. 2021), *C. elegans* (Gilbert et al. 2022), yeast (Elyashiv et al. 2010; Koufopanou et al. 2015; Huber et al. 2017), *Drosophila* (Campos et al. 2014; Castellano et al. 2015; Huang et al. 2021; Keightley and Eyre-Walker 2007; Huber et al. 2017), *Arabidopsis* (Moutinho, Trancoso, and Dutheil 2019; Huber et al. 2018), primates (Castellano et al. 2019; Galtier 2016; Ma et al. 2013; Hvilsom et al. 2012), and humans (Eyre-Walker and Keightley 2009; Boyko et al. 2008; Eyre-Walker, Woolfit, and Phelps 2006; Y. Li et al. 2010; Kim, Huber, and Lohmueller 2017; Huang et al. 2021). For humans, this approach has suggested that the DFE for nonsynonymous mutations in humans contains many nearly neutral mutations (e.g. 32% of mutations were estimated to have *s* > −10^−4^ by Kim et al. 2017). The proportion of strongly deleterious mutations (*s* < −10^−2^) has varied across studies, ranging from ∼25-35% of mutations (Kim, Huber, and Lohmueller 2017; Boyko et al. 2008; Eyre-Walker, Woolfit, and Phelps 2006). Importantly, recent studies using larger sample sizes have suggested that the proportion of very strongly deleterious mutations (*s* < −0.1) is very small (e.g., Kim et al. 2017 estimated ∼3%).

Studies of the DFE using genetic variation from natural populations summarized by the SFS have several limitations. First, inferences conducted to date typically assume that the effects of deleterious mutations are additive (though see Huber et al. 2018). This is largely due to computational convenience and the challenges in separately inferring dominance and the DFE using data from a single population. Second, the genetic variation data in a sample of a few hundred or fewer individuals from the population largely consists of weakly deleterious or neutral variation, as strongly deleterious mutations are unlikely to be segregating in samples of modest size (Kim, Huber, and Lohmueller 2017). Instead, the inferences made about the proportions of strongly deleterious mutations are informed by the absence of genetic variation expected under the particular mutation rate and demographic history and the functional form of the DFE assumed (Bank et al. 2014). Thus, the SFS-based methods to infer the DFE may not give a complete picture of the proportion of lethal and near-lethal mutations.

Given these limitations, several studies have recently suggested that there may be a much higher proportion of lethal or near-lethal new mutations in a variety of organisms. For example, Kardos et al. (2021) proposed a DFE where 5% of deleterious mutations are recessive lethal and ∼21% have *s* < -0.1. Similarly, Pérez-Pereira, Caballero, and García-Dorado (2022) proposed a DFE with that ∼4% of deleterious mutations are recessive lethal, and ∼41% have *s* < -0.1. Importantly, both of these DFEs were not directly estimated from data, but were instead designed largely to reflect results from mutation accumulation experiments (Kardos et al. 2021; Pérez-Pereira, Caballero, and García-Dorado 2022). Other recent studies have focused on “nearly lethal” mutations, or those mutations that are not expected to segregate in a sample of genetic variation. For example, Dukler et al. (2021) used genetic variation data from tens of thousands of human exomes and estimated that 22% of new mutations at 0-d sites are additive lethal or nearly lethal. Similarly, Galtier and Rousselle (2020) estimated that ∼60% of mutations are “lethal” based on genetic variation data from a diverse assemblage of animals, where lethal mutations are defined as having zero probability of being observed as polymorphic. In nearly all of these papers, it is argued that previous approaches to infer the DFE from genetic variation data have missed this large proportion of lethal and strongly deleterious mutations.

Here, we test the ability of one SFS-based approach, Fit*∂*a*∂*i (Kim, Huber, and Lohmueller 2017), to detect lethal mutations using simulated genetic variation data. We find that Fit*∂*a*∂*i is unable to accurately infer the simulated fraction of recessive lethals when performing inference under an additive model, as expected given the misspecification of the dominance model. However, we find that Fit*∂*a*∂*i can accurately estimate the simulated fraction of additive lethal mutations. Moreover, in both the recessive and additive case, we find that the presence of lethals does not greatly impact inference of the non-lethal portion of the DFE. Finally, as an alternative approach for quantifying the fraction of new mutations that are recessive lethal, we use mutation-selection-drift balance models in concert with estimates of segregating recessive lethals in humans and *Drosophila melanogaster*. These results demonstrate that a very small fraction of mutations (<1%) in *D. melanogaster* and humans are likely to be recessive lethal. Our work places limits on the proportion of mutations that are recessive lethal, which has implications for studying inbreeding depression in a variety of species.

## Results

### Can Fit *∂*a*∂*i infer the proportion of recessive lethals?

Due to their large impact on fitness, lethal mutations are typically fully recessive or nearly so (Simmons and Crow 1977), such that two copies of the allele must be present to have a detrimental impact on fitness. We focus first on the case of recessive lethals, specifically testing whether Fit*∂*a*∂*i can estimate the fraction of recessive lethals from simulated data where the true proportion of recessive lethal mutations is known. We used the forward-in-time simulator SLiM 3 (Haller and Messer 2019) to generate genetic variation datasets for use in Fit*∂*a*∂*i (see **Fig. S1**). We modeled a ∼30 Mb coding sequence with randomly-placed genic regions accumulating synonymous (neutral) and nonsynonymous (deleterious) mutations, some fraction of which were recessive lethal. Selection coefficients (*s*) for nonsynonymous mutations were drawn from a gamma DFE estimated from human genetic variation data (Kim, Huber, and Lohmueller 2017), with all non-lethal mutations assumed to be additive (dominance coefficient *h* = 0.5). To model different proportions of mutations that are recessive lethal, we assumed that either 0%, 1%, 5%, or 10% of new nonsynonymous mutations were recessive lethal (see Methods; **Fig. S1, Table S1**). These proportions roughly correspond to 0, 1.6, 7.9, and 15.8 recessive lethals per simulated diploid, respectively. For all simulations, we assumed a constant population size of *N*=10,000.

We began by examining the site frequency spectra (SFS) for models including recessive lethal mutations. In a sample of 1000 haploid individuals from the population, we found that recessive lethal mutations are segregating in the sample if the proportion of recessive lethals is at least 5% (**Fig. 1**).

**Figure 1.**
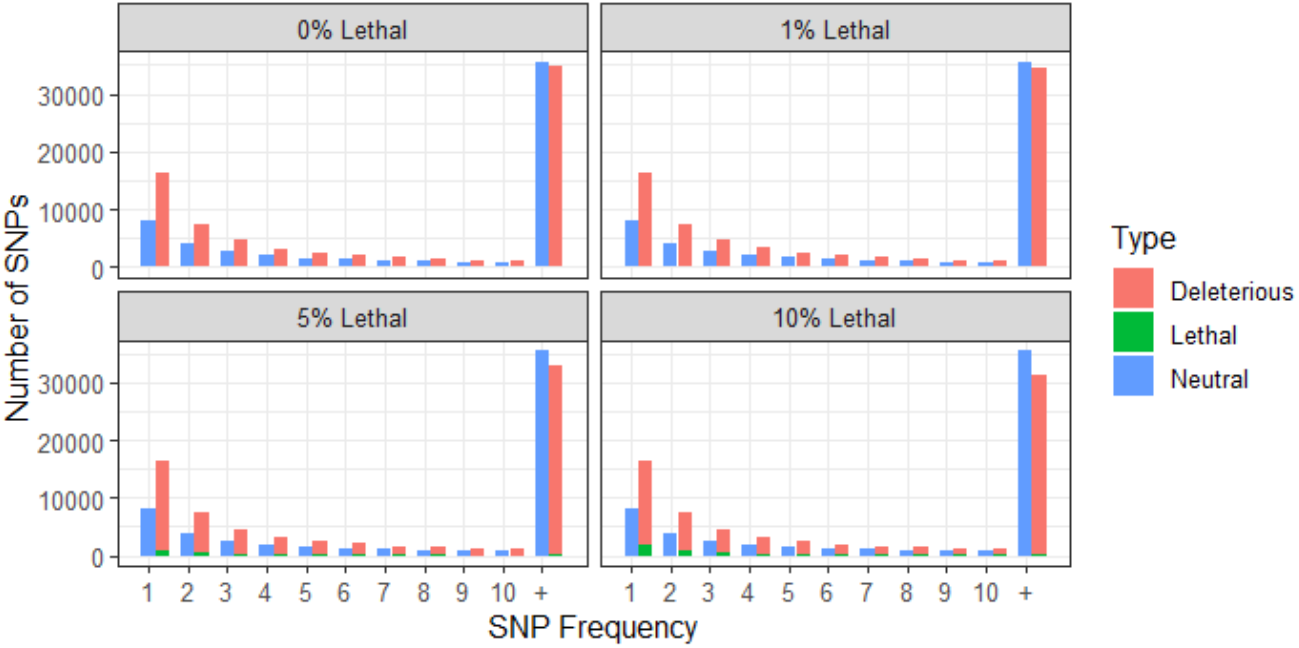
Unfolded site frequency spectra from simulated data with increments of recessive lethals. Colors represent the different mutation types (deleterious, neutral, and lethal). The X-axis depicts the frequency of the different mutation types (1 = singleton, 2 = doubleton, etc.), while the Y-axis depicts the number of mutations observed. SFS generated from 1000 sampled haploids.

To infer the DFE from these simulated data, we used Fit*∂*a*∂*i (Kim, Huber, and Lohmueller 2017). This software first relies on a class of putatively neutral mutations to estimate a demographic model (see **Table S1-2**), then, conditional on this demographic model and a given mutation rate, uses nonsynonymous variants to estimate the parameters of the distribution of fitness effects (DFE) of new nonsynonymous mutations.

We initially performed inference assuming that the DFE follows a gamma distribution and that all deleterious mutations have an additive effect on fitness (*h* = 0.5), exploring how varying levels of simulated recessive lethals impact the inference of the shape (*α*) and scale (*β*) parameters of this distribution (see Methods, **Figure 2A**). For presentation of these results, we discretized the DFE in categories defined as neutral (|*s*| = [0; 10^−5^), 0 < 2*Ns* < 0.1), nearly neutral (|*s*| = [10^−5^; 10^−4^), 0.1 < 2*Ns* < 1), slightly deleterious (|*s*| = [10^−4^; 10^−3^), 1 < 2*Ns* < 10), moderately deleterious (|*s*| = [10^−3^; 10^−2^), 10 < 2*Ns* <100), and strongly deleterious (|*s*| = [10^−2^, 1], 100 < 2*Ns* <10000).

**Figure 2.**
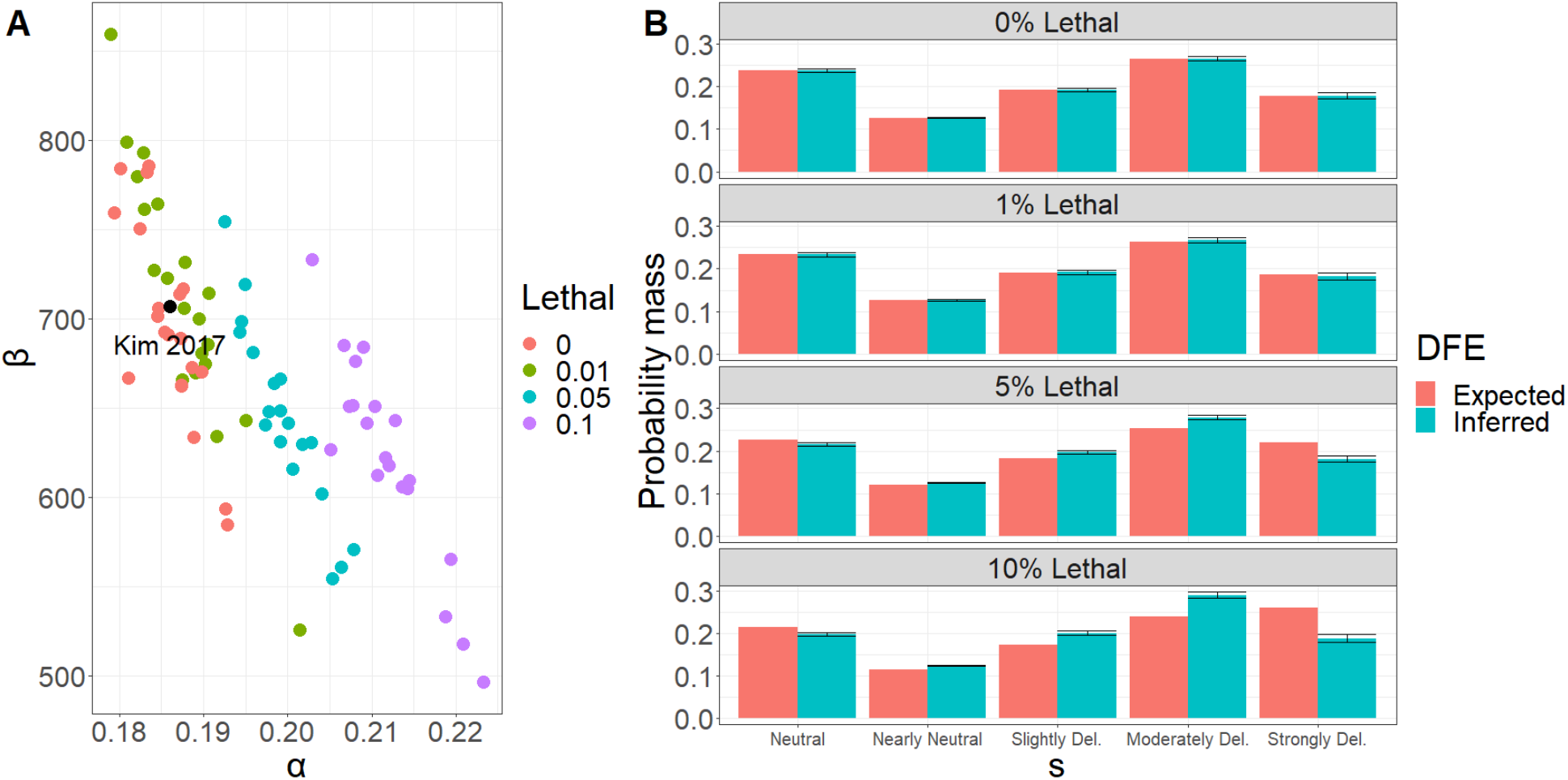
Parameters of the gamma distribution and the inferred DFE under different levels of recessive lethals A. Inference of the shape and scale parameters under a gamma DFE model from simulated data with different levels of recessive lethals using 1000 haploid individuals in each simulation. **B**. Expected versus inferred DFE for each lethal percentage under a gamma DFE model. Percentages are increments of recessive lethal mutations. Expected and inferred SFSs used to fit the model are presented in **Figure S2**. The DFE categories are defined as neutral (|*s*| = [0; 10^−5^)), nearly neutral (|*s*| = [10^−5^; 10^−4^)), slightly deleterious (|*s*| = [10^−4^; 10^−3^)), moderately deleterious (|*s*| = [10^−3^; 10^−2^)), and strongly deleterious (|*s*| = [10^−2^, 1]). Error bars correspond to the range of inferred proportions obtained across 20 simulation replicates.

The presence of recessive lethal mutations shifts the distributions of the estimates of the shape and scale parameters of the DFE (**Fig. 2A**). However, the introduction of recessive lethals has a relatively small impact on the inferred proportions of mutations in each category (**Figure 2B**), with the largest deviation observed in simulations with 10% of recessive lethals. Specifically, for a model where 10% of mutations are recessive lethal, we infer a notable deficit of strongly deleterious variation relative to the true proportions (**Figure 2B**). This deviation is likely due to underrepresentation of rare variants and overrepresentation of common variants relative to what is expected under a fully additive model (**Fig. S2**). In other words, recessive lethal mutations segregate at higher frequencies than expected under an additive model, and Fit*∂*a*∂*i therefore infers these mutations to have a *s* that is only moderately deleterious. Thus, when performing inference under an additive model, Fit*∂*a*∂*i cannot detect the presence of recessive lethals, as expected given the mis-specification of the dominance model.

Because sample size affects the ability to discriminate across different classes of mutations (Kim, Huber, and Lohmueller 2017), we performed the DFE inference under two smaller sample sizes of 10 and 100 haploids. As expected, the decrease in sample size increases the observed variance in DFE estimates across simulated datasets (**Fig. 3A**). The inferred fraction of moderately and strongly deleterious mutations is especially variable, independent of the proportions of lethals added to the simulation. This result is a consequence of mutations with |*s*| > 10^−3^ being unlikely to segregate in smaller sample sizes, thus impeding inference of the true proportion of moderately and strongly deleterious mutations. Moreover, we observe the counterintuitive result that the inferred DFEs with small sample sizes are closer to the true DFEs in the presence of high levels of recessive lethals, though with higher variance (**Fig. 3A**). This is likely because, with smaller sample sizes, recessive lethals do not segregate and impact DFE inference (**Fig. S3**), and the tail of the gamma DFE can then be extrapolated from the more weakly deleterious mutations that are segregating, provided that the functional form of the DFE is known.

**Figure 3.**
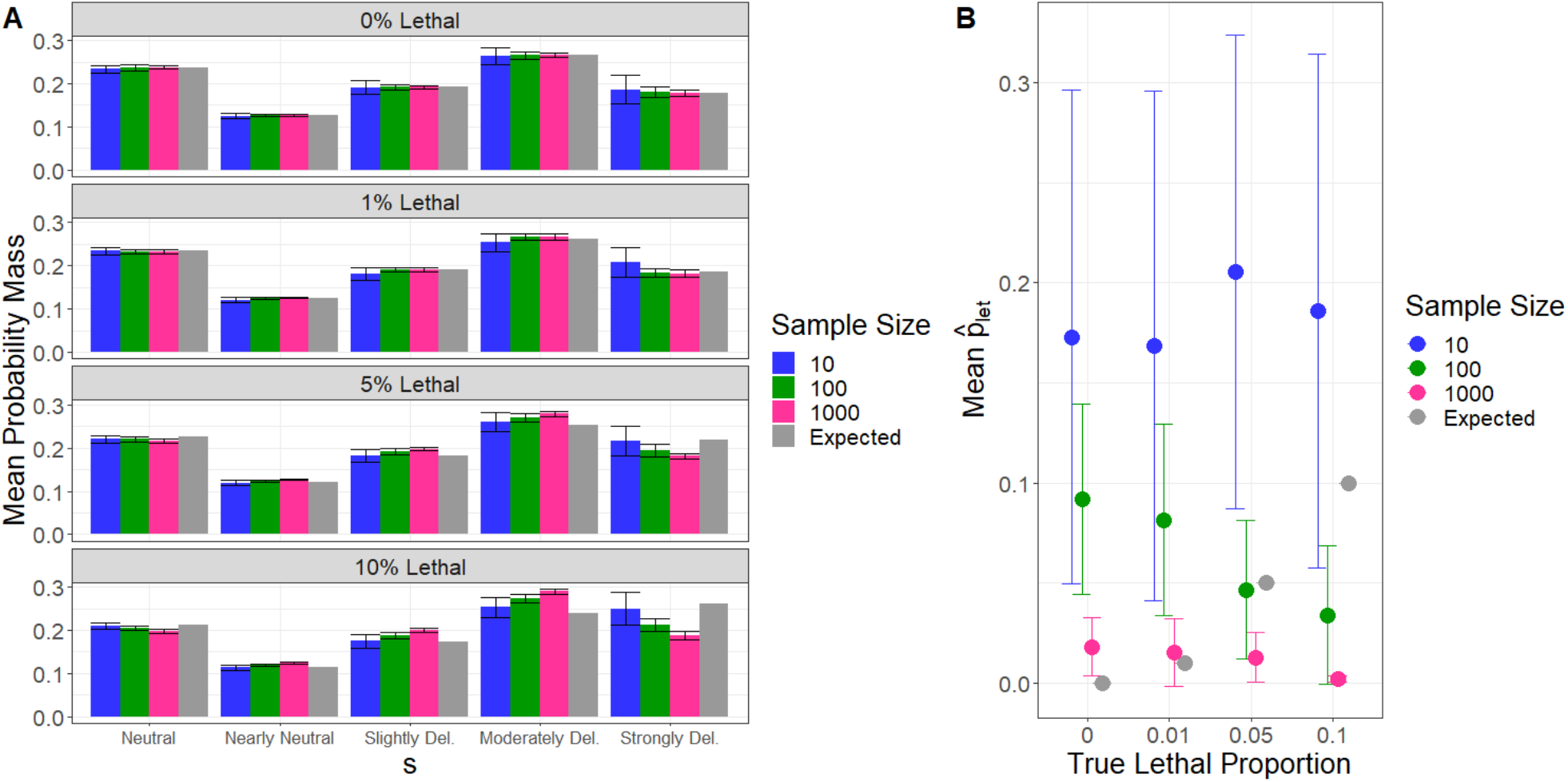
Inferred DFE and proportion of recessive lethals 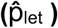 depending on the sample size and on the true portion of recessive lethals mutations simulated. **A**. The mean probability mass obtained with the gamma parameters for each DFE category. The DFE categories are defined as neutral (|*s*| = [0; 10^−5^)), nearly neutral (|*s*| = [10^−5^; 10^−4^)), slightly deleterious (|*s*| = [10^−4^; 10^−3^)), moderately deleterious (|*s*| = [10^−3^; 10^−2^)), and strongly deleterious (|*s*| = [10^−2^, 1]). Error bars correspond to the range of inferred proportions obtained across 20 simulation replicates. **B**. The mean proportion of lethals inferred 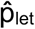 with the Gamma+Let model and different sample sizes. The gray dots correspond to the expected true proportion of lethals (p_let_). Error bars correspond to the range of values of 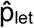 inferred from 20 simulation replicates.

Next, we attempted to fit more complex distributions of fitness effects to determine whether adding a point mass of lethal mutations may improve inference of the fraction of recessive lethals. Specifically, we evaluated whether DFEs allowing for a mixture distribution of a point mass of neutral (Neu+Gamma model), a point mass of lethal (Gamma + Let), and a point mass of neutral and lethal (Neu+Gamma+Let) distributed new mutations, would fit the simulated nonsynonymous SFS better under the different increments of lethals than the Gamma DFE (**Table 1**). Throughout, we refer to our estimate of the fraction of new mutations that are lethal as 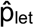.

**Table 1.** Maximum likelihood estimates (MLEs) assuming different distribution of fitness effects when simulating varying proportions of recessive lethals. These results are reported assuming a nonsynonymous to synonymous sequence length ratio (LNS/LS) of 2.31, mutation rate (μ) = 1.5×10^−8^, and 1000 simulated haploid individuals. For each DFE, 20 replicates were conducted and average estimates are reported.

| Lethal Prop. | DFE | Average Parameter MLEs | Log-likelihood | AIC |
| --- | --- | --- | --- | --- |
| 0% | Gamma | $\alpha = 0.186, \beta = 696.06$ | -2811.696634 | 5627.393 |
| | Neu+Gamma | $\alpha = 0.2, \beta = 643.483, \hat{p}_{neu} = 0.033$ | -2812.326619 | 5630.653 |
| | Gamma+Let | $\alpha = 0.188, \beta = 597.446, \hat{p}_{let} = \mathbf{0.018}$ | -2809.970874 | **5625.942** |
| | Neu+Gamma+Let | $\alpha = 0.209, \beta = 569.175, \hat{p}_{neu} = 0.045, \hat{p}_{let} = 0.011$ | -2811.503446 | 5631.007 |
| 1% | Gamma | $\alpha = 0.188, \beta = 711.919$ | -2808.123191 | 5620.246 |
| | Neu+Gamma | $\alpha = 0.206, \beta = 643.71, \hat{p}_{neu} = 0.041$ | -2808.389979 | 5622.78 |
| | Gamma+Let | $\alpha = 0.189, \beta = 631.715, \hat{p}_{let} = \mathbf{0.015}$ | -2806.116316 | **5618.233** |
| | Neu+Gamma+Let | $\alpha = 0.25, \beta = 533.182, \hat{p}_{neu} = 0.089, \hat{p}_{let} = 0.011$ | -2826.773773 | 5661.548 |
| 5% | Gamma | $\alpha = 0.2, \beta = 644.815$ | -2799.75543 | 5603.511 |
| | Neu+Gamma | $\alpha = 0.244, \beta = 517.21, \hat{p}_{neu} = 0.076$ | -2797.735431 | **5601.471** |
| | Gamma+Let | $\alpha = 0.201, \beta = 588.789, \hat{p}_{let} = 0.013$ | -2799.140014 | 5604.28 |
| | Neu+Gamma+Let | $\alpha = 0.245, \beta = 479.858, \hat{p}_{neu} = 0.074, \hat{p}_{let} = 0.012$ | -2797.715242 | 5603.43 |
| 10% | Gamma | $\alpha = 0.212, \beta = 621.492$ | -2786.548426 | 5577.097 |
| | Neu+Gamma | $\alpha = 0.34, \beta = 365.471, \hat{p}_{neu} = 0.145$ | -2771.432743 | **5548.865** |
| | Gamma+Let | $\alpha = 0.212, \beta = 615.501, \hat{p}_{\text{let}} = 0.002$ | -2786.796788 | 5579.594 |
| | Neu+Gamma+Let | $\alpha = 0.32, \beta = 390.033, \hat{p}_{\text{neu}} = 0.114, \hat{p}_{\text{let}} = 0.012$ | -2775.605873 | 5559.212 |

For the case of 0% or 1% simulated recessive lethals, we find that the Gamma+Let model fits the data best. Additionally, in the 0% lethals case, we estimate 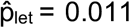, and in the 1% lethals case, we estimate 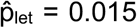 (**Table 1**). When simulating higher fractions of recessive lethals (5% and 10%), we find that the Neu+Gamma model fits the data best. Moreover, estimates of 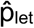 under a Gamma+Let model are much lower than the simulated values, remaining in all cases close to zero (**Table 1**). Sample size does not affect 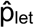 accuracy: the true p_let_ is largely mis-inferred in smaller sample sizes, and the inclusion of 1000 haploids does not improve accuracy (**Fig. 3B**).

Finally, to evaluate Fit*∂*a*∂*i’s performance under a Gamma+Let model further, we conducted log-likelihood profiling by considering a grid of values for 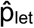. For each value of p_let_, we then re-optimized the other DFE parameters (*α* and *β*). We then evaluated the difference in log-likelihood between the mean maximum likelihood estimation (MLE) across replicates and the log-likelihood under a given true p_let_. Here, we again find that Fit*∂*a*∂*i cannot accurately estimate the true fraction of simulated recessive lethals, and performs especially poorly when the DFE contains a high proportion of recessive lethal mutations (5% and 10%; **Fig. S4**). Thus, we conclude that we have limited power to infer the correct fraction of lethal mutations when lethals are completely recessive.

### Can Fit *∂*a *∂*i infer the proportion of additive lethals?

Our inability to accurately detect lethal mutations when they are fully recessive is not surprising, given that Fit*∂*a*∂*i’s assumes, by default, that all mutations have additive effects on fitness (Kim, Huber, and Lohmueller 2017). To determine how much our inferences are improved when the data are generated under the dominance model used for inference, we next performed simulations where lethal mutations were assumed to be additive.

As expected, additive lethals do not segregate in the simulated datasets of sample sizes of 1000 haploid individuals (**Fig. 4**), given that they are immediately removed by purifying selection. When estimating the DFE under a gamma model, we find that the inferred DFEs are much closer to the true DFEs when compared to results for recessive lethals (**Fig. 5**). Specifically, the *α* parameters estimated under the different increments of additive lethals are less variable than in the recessive case (*α* range between 0.174 and 0.186 for the additive model, and between 0.186 and 0.212 for the recessive case). However, the *β* parameter varied more in the additive scenario (*β* ranged [702,1814] for the additive case and ranged [621,712] for the recessive case) (**Fig. 5A**).

**Figure 4.**
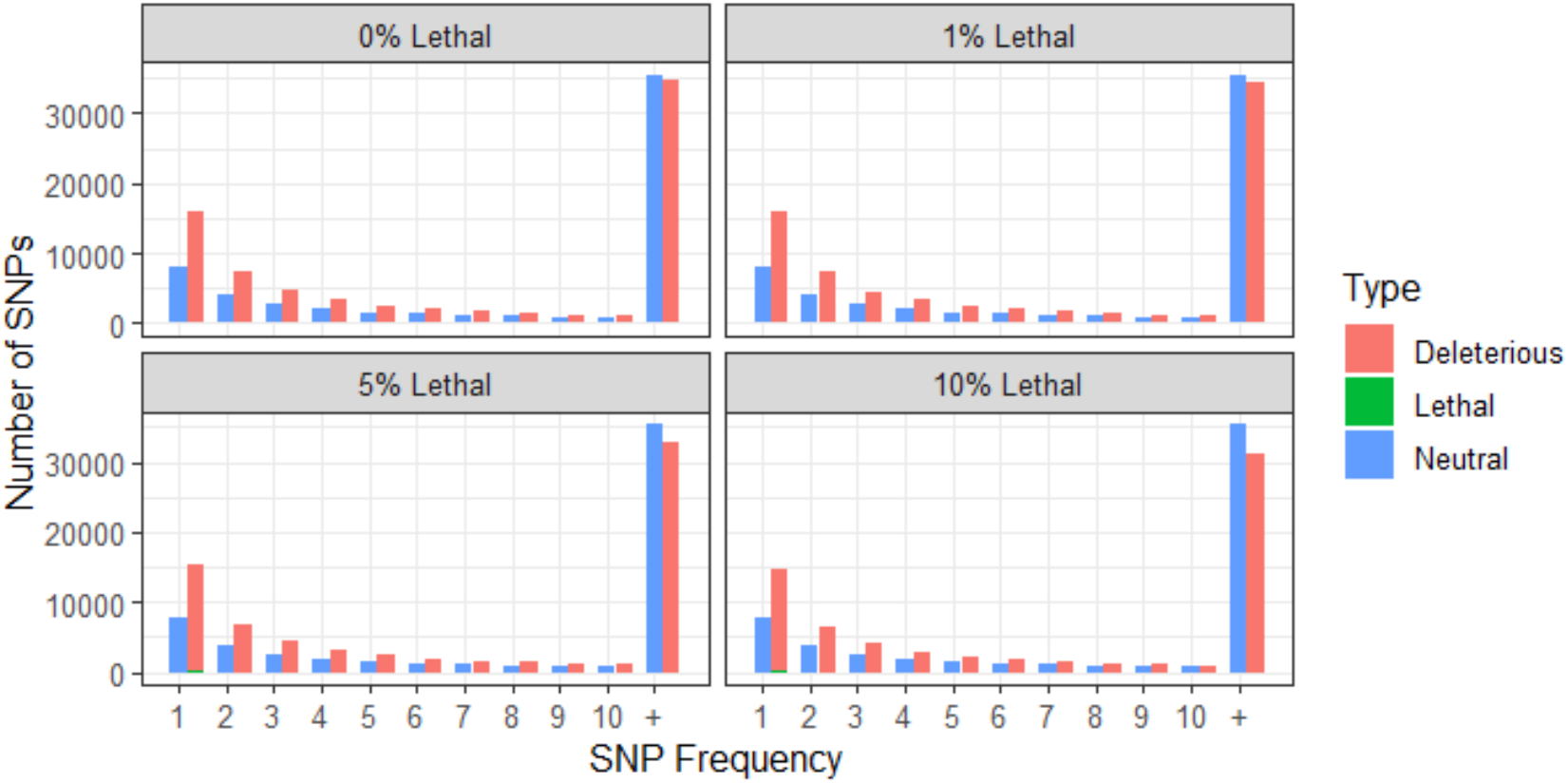
Unfolded site frequency spectra from simulated data with increments of additive lethals in a population size of 1000 haploid individuals. Colors represent the different mutation types (deleterious, neutral, and lethal). The X-axis depicts the frequency of the different mutation types (1 = singleton, 2 = doubleton, etc.), while the Y-axis depicts the number of mutations observed. Note that no lethals were observed segregating, as expected when lethals are additive. SFS generated from 1000 sampled haploids.

**Figure 5.**
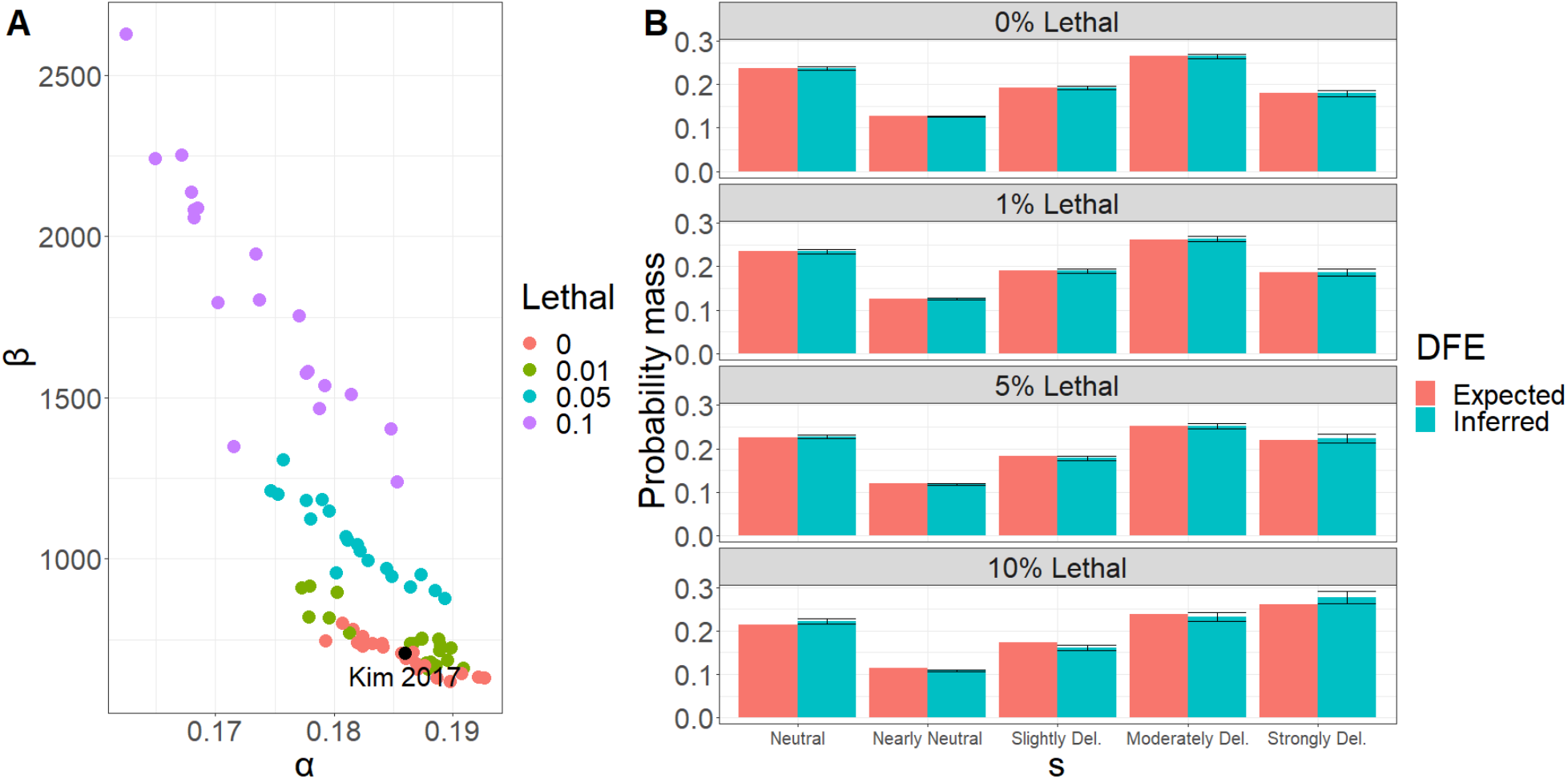
Parameters of the gamma distribution and the inferred DFE under different levels of additive lethals. A. Inference of the shape and scale parameters under a gamma DFE model from simulated data with different levels of additive lethals in a population size of 1000 haploid individuals. **B**. Expected versus inferred DFE for each lethal percentage under a gamma DFE, percentages are increments of additive lethal mutations. Expected and inferred SFSs used to fit the model are presented in **Figure S2**. The DFE categories are defined as neutral (|*s*| = [0; 10^−5^)), nearly neutral (|*s*| = [10^−5^; 10^−4^)), slightly deleterious (|*s*| = [10^−4^; 10^−3^)), moderately deleterious (|*s*| = [10^−3^; 10^−2^)), and strongly deleterious (|*s*| = [10^−2^, 1]. Error bars correspond to the range of inferred proportions obtained across 20 simulation replicates.

When fitting more complex models of the DFE to this data, we consistently find Gamma+Let to be the best-fitting model (**Table 2**). Moreover, our estimates of 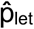 are generally close to the true simulation fraction of lethals. Specifically, when assuming lethal levels of 1%, we Infer 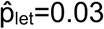, when assuming lethal levels of 5%, we infer 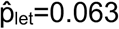, and when assuming lethal levels of 10%, we infer 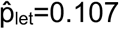 (**Table 2**). As previously observed for recessive lethals, the sample size does not affect the inference of the expected non-lethal portion of the DFE (**Fig. S5A**), but sample size does improve 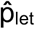 estimation (**Fig. S5B**).

**Table 2.** Maximum likelihood estimates (MLEs) assuming different distribution of fitness effects when simulating varying proportions of additive lethals. These results are reported assuming a nonsynonymous to synonymous sequence length ratio (LNS/LS) of 2.31, mutation rate (μ) = 1.5×10^−8^, and 1000 simulated haploid individuals. For each DFE, 20 replicates were conducted and average estimates are reported.

| Lethal Prop. | DFE | Average Parameter MLEs | Log-likelihood | AIC |
| --- | --- | --- | --- | --- |
| 0% | Gamma | $\alpha = 0.186, \beta = 696.06$ | -2811.696634 | 5627.393 |
| | Neu+Gamma | $\alpha = 0.2, \beta = 643.483, \hat{p}_{\text{neu}} = 0.033$ | -2812.326619 | 5630.653 |
| | Gamma+Let | $\alpha = 0.188, \beta = 597.446, \hat{p}_{\text{let}} = \mathbf{0.018}$ | -2809.970874 | **5625.942** |
| | Neu+Gamma+Let | $\alpha = 0.209, \beta = 569.175, \hat{p}_{\text{neu}} = 0.045, \hat{p}_{\text{let}} = 0.011$ | -2811.503446 | 5631.007 |
| 1% | Gamma | $\alpha = 0.186, \beta = 753.876$ | -2820.718696 | 5645.437 |
| | Neu+Gamma | $\alpha = 0.2, \beta = 698.046, \hat{p}_{\text{neu}} = 0.032$ | -2821.248792 | 5648.498 |
| | Gamma+Let | $\alpha = 0.189, \beta = 585.229, \hat{p}_{\text{let}} = \mathbf{0.03}$ | -2815.502511 | **5637.005** |
| | Neu+Gamma+Let | $\alpha = 0.214, \beta = 562.23, \hat{p}_{\text{neu}} = 0.054, \hat{p}_{\text{let}} = 0.021$ | -2817.933003 | 5643.866 |
| 5% | Gamma | $\alpha = 0.182, \beta = 1056.832$ | -2820.940223 | 5645.88 |
| | Neu+Gamma | $\alpha = 0.199, \beta = 940.111, \hat{p}_{\text{neu}} = 0.038$ | -2821.863918 | 5649.728 |
| | Gamma+Let | $\alpha = 0.188, \beta = 614.993, \hat{p}_{\text{let}} = \mathbf{0.063}$ | -2801.582183 | **5609.164** |
| | Neu+Gamma+Let | $\alpha = 0.219, \beta = 604.409, \hat{p}_{\text{neu}} = 0.06, \hat{p}_{\text{let}} = 0.05$ | -2806.160594 | 5620.321 |
| 10% | Gamma | $\alpha = 0.174, \beta = 1813.546$ | -2808.35107 | 5620.702 |
| | Neu+Gamma | $\alpha = 0.188, \beta = 1615.745, \hat{p}_{\text{neu}} = 0.03$ | -2810.254238 | 5626.508 |
| | Gamma+Let | $\alpha = 0.187, \beta = 639.097, \hat{p}_{\text{let}} = 0.107$ | -2754.489466 | **5514.979** |
| | Neu+Gamma+Let | $\alpha = 0.215, \beta = 581.44, \hat{p}_{\text{neu}} = 0.052, \hat{p}_{\text{let}} = 0.106$ | -2756.470155 | 5520.94 |

Finally, we again performed log-likelihood profiling to more closely examine Fit*∂*a*∂*i’s performance under different levels of simulated additive lethals. Here, we find that estimates of 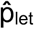 are consistently close to the true p_let_ (**Fig. 6**). However, the average 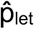 for 0% and 1% lethals are over-estimates, likely due to the true parameter being near the boundary of the parameter space. In conclusion, these results suggest that Fit*∂*a*∂*i performs reasonably well at inferring the proportion of lethal mutations when they are assumed to be additive and when the functional form of DFE is properly specified.

**Figure 6.**
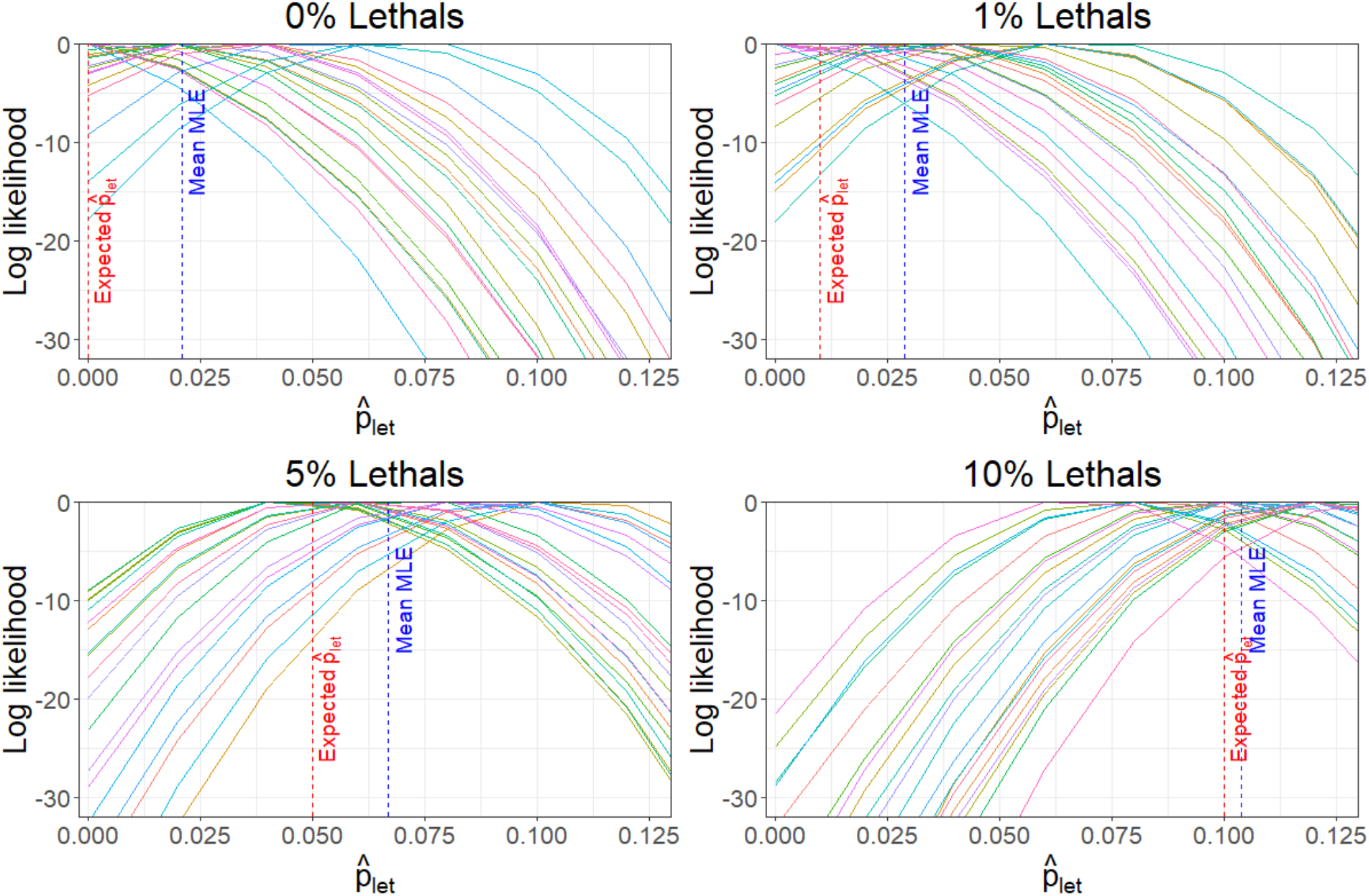
Evaluating the performance of Fit*∂*a*∂*i under a Gamma + lethal DFE when simulating additive lethals. The x-axis indicates the proportion of additive lethals, and y-axis denotes the log-likelihood for each value of *p*_*le*t_. Each line corresponds to a simulation replicate under different increments of additive lethals in a population size of 1000 haploids. Red dashed lines correspond to the expected (true) lethal proportion under each lethal simulated class, while blue dashed lines represent the mean maximum likelihood estimate across 20 independent replicates for each lethal class.

#### Using mutation-selection-drift balance to estimate the fraction of recessive lethals

Our results demonstrate that Fit*∂*a*∂*i is limited in its ability to estimate the fraction of new mutations that are recessive lethal, primarily due to violating the assumption of additivity. Although it is possible to conduct inference of recessive lethals under a recessive model, we have extremely limited information on the broader distribution of dominance coefficients in humans, which may confound inferences. Thus, we instead employ a different approach for quantifying the fraction of new mutations that are recessive lethal using models of mutation-selection-drift balance. Specifically, our aim was to use existing estimates of mutation rates, coding sequence length, effective population size (*Ne*) and segregating recessive lethal load in humans and *Drosophila melanogaster* to determine what fraction of new nonsynonymous mutations are recessive lethal. This approach relies on the result from (Nei 1968) relating the allele frequency of a recessive lethal mutation (*q*) to the mutation rate and effective population size:

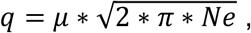

where μ is the mutation rate, *Ne* is the diploid effective population size, and *π* is the mathematical constant. Importantly, this model assumes that 2 * *Ne* * *μ* << 1 and that populations are at demographic equilibrium (i.e., constant population size). Thus, we explore a range of effective population sizes for each species, informed by simulations under a human demographic model (see **Supplementary Information**).

Our aim with this model was to determine what fraction of new nonsynonymous mutations would need to be recessive lethal to explain empirical estimates of the number of segregating recessive lethals per diploid in humans and *D. melanogaster*. As this model predicts an allele frequency for lethal mutations for a single site, our approach consists of multiplying this allele frequency by the nonsynonymous coding length for each species (**Table 3**). We then assessed the proportion of mutations that would need to be recessive lethal to explain observed numbers of segregating lethal mutations. Given that our analysis assumes that recessive lethals can only arise as nonsynonymous mutations, whereas estimates of segregating lethals are genome-wide, results from this approach should therefore be interpreted as upper bounds.

**Table 3:** Summary of parameters used in mutation-selection-drift balance analyses. The assumed mutation rate (*μ*), effective population size (Ne), total coding length in base-pairs (L), nonsynonymous to synonymous ratio (NS:SYN ratio) is shown for each species. Also shown is the range of segregating recessive lethals per diploid estimated in humans and *D. melanogaster*. See Methods for references and further detail.

| Parameters | Humans | *D. melanogaster* |
| --- | --- | --- |
| $\mu$ | 1.50e-08 | 3.00e-09 |
| $Ne$ | 20,000-60,000 | 5e5-5e6 |
| $L$ | 30,150,000 | 22,005,320 |
| NS:SYN ratio | 2.31:1 | 2.85:1 |
| # segregating lethals | 0.6-1.6 | 1-3 |

We first explored predictions of this model using mutation rate and coding sequence length estimates for humans (see Methods; **Table 3**). When evaluating model predictions for a range of effective population sizes of *Ne* = {20000, 30000, 60000}, we consistently find that empirical estimates of the number of recessive lethals per diploid in humans can be explained by a small proportion of new mutations being recessive lethal (**Fig. 7**). Specifically, available evidence suggests that the average number of segregating recessive lethal mutations in humans is roughly in the range of 0.6 to 1.6 per diploid (Gao et al. 2015; Narasimhan et al. 2016), and analytical predictions with <∼0.5% of new nonsynonymous mutations being recessive lethal appear to approximate this range, regardless of the assumed effective population size. By contrast, analytical predictions with ∼5% of new mutations being recessive lethal, a value that has recently been proposed (Kardos et al. 2021; Pérez-Pereira, Caballero, and García-Dorado 2022), suggest ∼10-20 recessive lethal mutations per diploid, well outside of empirical observations.

**Figure 7:**
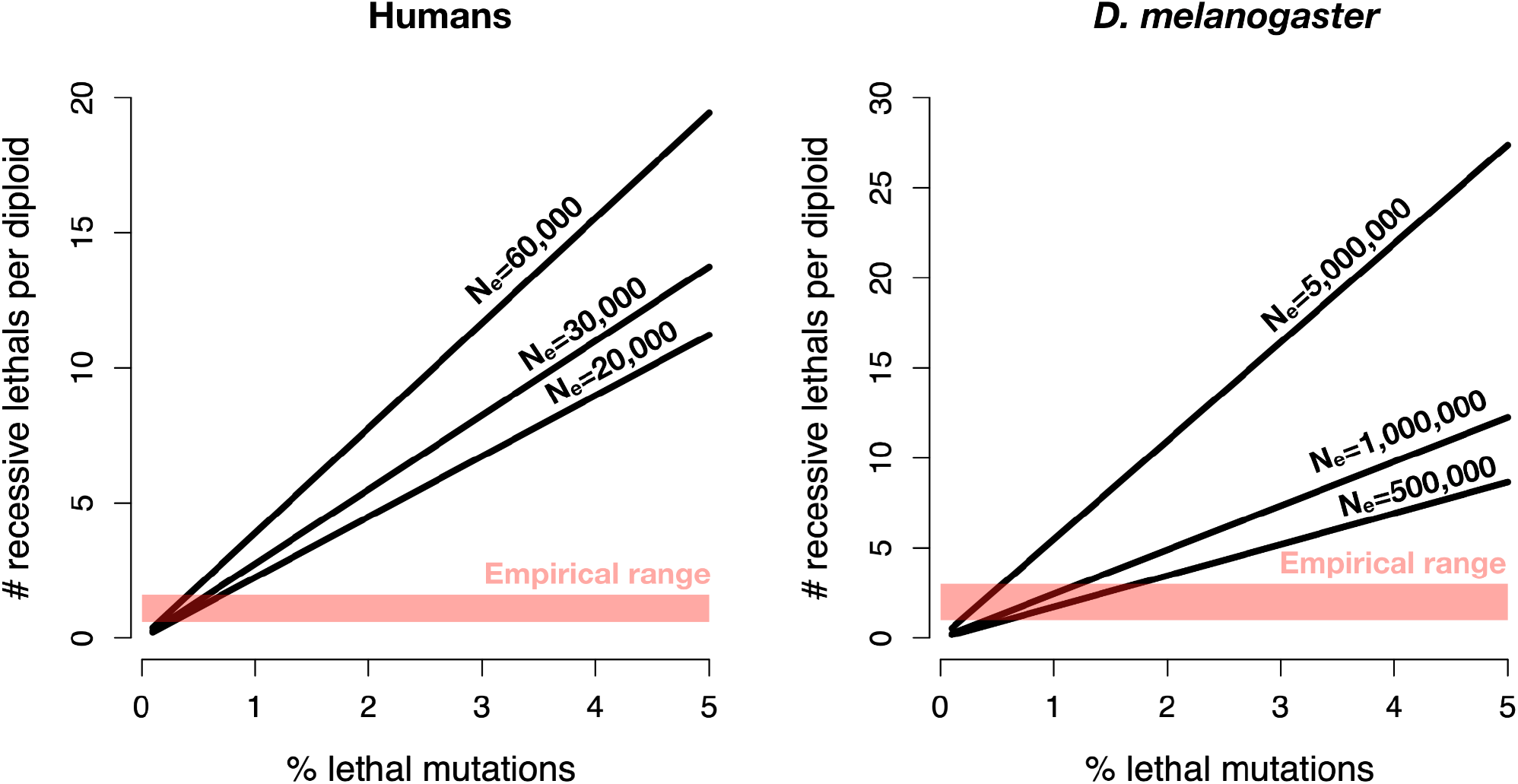
Relationship between the percent of new mutations that are recessive lethal and the predicted number of segregating recessive lethals per diploid under mutation-selection-drift balance for humans and *Drosophila melanogaster* under varying effective population sizes. X-axis denotes the percent of new nonsynonymous mutations that are recessive lethal and Y-axis denotes the resulting number of segregating recessive lethals per diploid. Red shading indicates the range of empirical estimates of segregating recessive lethals for each species. Note that, in both cases, the overlap between the empirical estimates and model prediction suggests that <1% of new nonsynonymous mutations are likely to be recessive lethal.

We next explored predictions of this model using mutation rate and coding sequence length estimates for *Drosophila melanogaster* (see Methods, **Table 3**). We computed the expected number of recessive lethals per diploid assuming three different effective population sizes *Ne* = {500000, 1000000, 5000000}, reflecting the range of estimated effective population sizes in the literature (Duchen et al. 2013; Sheehan and Song 2016; Huber et al. 2017; H. Li and Stephan 2006). Here, we again find that empirical estimates of segregating recessive lethals can be explained by a small fraction of new mutations being recessive lethal (**Fig. 7**). Specifically, available evidence suggests ∼1-3 recessive lethals per diploid in *D. melanogaster* (Simmons and Crow 1977; McCune et al. 2002; Lynch, Walsh, and Others 1998), which can be explained in our model by <∼1% of new nonsynonymous mutations being recessive lethal, though results are dependent on the assumed effective population size (**Fig. 7**). Importantly, a model where 5% of nonsynonymous mutations are assumed to be recessive lethal predicts ∼9-27 recessive lethals per diploid, well beyond the empirical range for *D. melanogaster*.

In summary, these results for humans and *D. melanogaster* consistently find that a small fraction (<∼1%) of new nonsynonymous mutations are likely to be recessive lethal. Although this model makes a number of simplifying assumptions, results are strikingly consistent across species and appear to be relatively robust to recent exponential growth (see **SI**). Moreover, in both cases, we find that a proposed fraction of 4-5% of new mutations being recessive lethal (Kardos et al. 2021; Pérez-Pereira, Caballero, and García-Dorado 2022) predicts levels of segregating lethal variation that are well outside of empirical estimates. Thus, our results place an upper bound on the recessive lethal portion of the DFE, suggesting that no more than ∼1% of new nonsynonymous mutations are recessive lethal.

## Discussion

In this study, we have explored different approaches for estimating the proportion of new mutations that are lethal. Our results demonstrate the limitations of SFS-based methods for detecting recessive lethal mutations, while also providing an estimate of the fraction of recessive lethal mutations based on models of mutation-selection-drift balance. Specifically, we show using simulations that a commonly-used SFS-based method for DFE inference, Fit*∂*a*∂*i (Kim, Huber, and Lohmueller 2017), cannot accurately quantify the proportion of new mutations that are recessive lethal (**Fig 1, Table 1, Fig S4**). This result is not surprising, given that Fit*∂*a*∂*i assumes all mutations are additive by default, whereas our simulations assumed lethal mutations to be fully recessive (*h*=0.0) and all other non-lethal deleterious mutations to be additive (*h*=0.5). However, despite our inability to correctly infer the fraction of recessive lethals, we demonstrate that the presence of recessive lethals has a relatively minimal impact on inference of the deleterious non-lethal portion of the DFE, especially at low levels of recessive lethals (**Fig. 2**). Thus, to the extent that recessive lethals do comprise a small portion of the DFE (**Fig. 7**), and that the deleterious non-lethal portion of the DFE truly is additive and gamma distributed, our results suggest that existing estimates of this portion of the DFE are robust to the presence of lethals.

In reality, there is likely to be an inverse relationship between the selection coefficient of a mutation and its dominance coefficient, such that strongly deleterious mutations are highly recessive and nearly neutral mutations are closer to additive (Huber et al. 2018; Agrawal and Whitlock 2011). Our analysis largely ignores this complexity by assuming mutations to be either fully additive or fully recessive. Future work on refining DFE estimates in humans and other species should incorporate this relationship between *h* and *s*, though doing so will require independent information on *h* and *s*, something that represents a major challenge (Huber et al. 2018; Fuller et al. 2019). However, recent theoretical work suggests that combining information from the SFS along with linkage disequilibrium may provide sufficient power for obtaining a joint distribution of *s* and *h* (Ragsdale 2021).

In contrast to our results for recessive lethals, we find that Fit*∂*a*∂*i can accurately infer the fraction of lethal mutations when they are assumed to be additive. This result further emphasizes that the violation of dominance assumptions is the main factor hindering the estimation of recessive lethals. However, the extent to which additive lethals actually exist, and if so, at what abundance, remains unclear. For example, a recent analysis of genetic variation data from ten animal taxa concluded that as much as 65% of new nonsynonymous mutations may be additive “lethal” (Galtier and Rousselle 2020). However, this paper defines lethals as mutations that “cannot contribute to observable polymorphism”, a definition that differs in meaningful ways from an actual lethal mutation and that depends on sample size. Similarly, Dukler et al. (2021) attempted to quantify the number of new “lethal or nearly lethal” mutations by examining sites depleted of variation in ∼72,000 human genomes, obtaining an estimate of ∼0.3–0.4 *de novo* lethal or nearly lethal mutations per potential human zygote. Here again, the extent to which sites depleted of genetic variation reflects the presence of lethal mutations is not entirely clear. Demonstrating this, Agarwal and Przeworski (2021) used a similar approach by examining invariant methylated CpG sites in a sample of 390,000 human genomes, concluding that an additive mutation with a selection coefficient as small as *s* = 1*e* − 3 is sufficient to result in a lack of variation even in the large sample sizes they used. Thus, approaches for detecting lethal mutations based on genetic variation data may have inherent limitations not only in disentangling *s* from *h*, but also in separating lethal mutations from mutations that are strongly deleterious though far from lethal (*s* on the order of -0.001 to -0.1).

Given the above challenges in quantifying recessive lethals using genetic variation data, we instead sought to obtain an estimate of the recessive lethal portion of the DFE by employing models of mutation-selection-drift balance. This approach leverages direct estimates of segregating recessive lethals in humans and *D. melanogaster* to provide an estimate of the fraction of new mutations that are recessive lethal. For both humans and *Drosophila*, we find that the observed number of segregating recessive lethals can be explained by <∼1% of new nonsynonymous mutations being recessive lethal (**Fig. 7**). This result implies a *de novo* mutation rate of ∼0.003 recessive lethal mutations per diploid human (assuming a lethal mutation rate of 0.5%) and ∼0.001 recessive lethal mutations per diploid fly (assuming a lethal mutation rate of 1%). Notably, when assuming 4-5% of new nonsynonymous mutations to be recessive lethal, as recently suggested (Kardos et al. 2021; Pérez-Pereira, Caballero, and García-Dorado 2022), these models predict levels of segregating recessive lethals that are well above those observed empirically (**Fig. 7**). Thus, these results suggest that we can reject the proposal that this number is as high as 4-5% (Kardos et al. 2021; Pérez-Pereira, Caballero, and García-Dorado 2022).

Our approach based on mutation-selection-drift balance makes a number of simplistic assumptions, which could impact our results. First, mutation-selection-drift balance models assume that populations are at equilibrium, whereas humans and *D. melanogaster* are known to have experienced recent exponential growth (Gravel et al. 2011; Tennessen et al. 2012; Duchen et al. 2013). To explore the impact of this assumption, we ran simulations under a human demographic model, finding that increases in recessive lethal mutation frequencies lag somewhat behind exponential growth. Consequently, the numbers of recessive lethal mutations roughly reflect the equilibrium value for a population with *Ne* on the order of 20,000 to 30,000 (see **SI, Fig. S6**), in agreement with results from Amorim et al. (2017). Although we were unable to conduct such simulations for *D. melanogaster* due to computational limitations, we instead projected recessive lethal values for a wide range of effective population sizes from 5e5 to 5e6 (**Fig. 7**), a range that encompassas existing *Ne* estimates for *D. melanogaster* (Duchen et al. 2013; Sheehan and Song 2016; Huber et al. 2017; H. Li and Stephan 2006). In the event that the impact of recent exponential growth in humans and *D. melanogaster* remains underestimated by our analysis, this would have the effect of only further reducing our estimate of the fraction of new mutations that are recessive lethal.

Another important assumption made by our analysis is that recessive lethal mutations can only arise as nonsynonymous mutations. This appears to be a reasonable assumption as a first approximation, especially in light of evidence that recessive lethal load does not appear to be influenced by genome size (McCune et al. 2002). However, it remains possible that lethal mutations can arise in other regions of the genome, such as conserved non-coding regions. As with our demographic assumptions, however, relaxing this assumption to include a greater length of sequence in our analysis would only further decrease the estimated fraction of new mutations that are recessive lethal. Finally, numerous other factors could contribute to recessive lethals segregating at higher levels than predicted by a simple model of mutation-selection-drift balance, such as epistasis, overdominance, and incomplete penetrance (Ballinger and Noor 2018; Amorim et al. 2017). Thus, these factors suggest that our estimate of ∼0.5% of new mutations being recessive lethal in humans likely represents an upper bound, implying that recessive lethals constitute a very small, though nevertheless important, portion of the DFE.

## Materials and methods

### Simulations

We made extensive use of simulations to (1) understand the impact of lethal mutations on the SFS and to (2) assess Fit*∂*a*∂*i’s performance (Kim, Huber, and Lohmueller 2017) in inferring the distribution of fitness effects (DFE) including lethals. We used the forward-in-time simulation software SLiM 3 (Haller and Messer 2019) to simulate genetic variation data using parameters that approximate those of a human genome. Specifically, we generated 30 Mb of coding sequence by simulating 30 chromosomal segments each of length 1Mb, with randomly placed coding regions comprising ∼1.5% of each segment (or ∼1.5 Mb coding sequence across the 30 segments). From each simulation, we computed both deleterious (non-lethal and lethal) and neutral site frequency spectra (SFS) from sample sizes of 10, 100 and 1000 haploids independently. The SFS were computed for each chromosomal segment separately and summed across the 30 segments. The resulting neutral and deleterious SFS were used for downstream demographic and the DFE inference, respectively.

Within each coding region, we modeled deleterious (nonsynonymous) and neutral (synonymous) mutations occurring at a ratio of 2.31:1 (Huber et al. 2017). To maintain computational efficiency, we did not model neutral mutations occurring in non-coding regions. We modeled a bimodal DFE for deleterious mutations, with non-lethal mutations arising from a gamma distribution (described by its shape (*α*) and scale (*β*) parameters) and lethal mutations arising with a fixed selection coefficient *s* = −1.0. The parameters of the gamma distribution were set based on estimates from the 1000G European population consisting of a mean *s* = -0.0131 and shape parameter of 0.186 (Kim, Huber, and Lohmueller 2017). The expected mean selection coefficient (*s* = -0.0131) was obtained by the shape (−0.186), scale (875), and ancestral population size (Na, 12,378) described in Kim, Huber, and Lohmueller (2017) as *E*[*s*] = (*scale* * *shape*)/2*Na*. We assumed varying levels of lethal mutations of 0%, 1%, 5%, and 10%. For example, in the case of 5% lethals, 95% of nonsynonymous mutations were drawn from the gamma distribution and 5% were set to have a selection coefficient *s* = −1.0 (see **Table S1**). All mutations arising from the gamma distribution were assumed to be additive (*h* = 0.5), whereas lethal mutations were assumed either to be additive or fully recessive (*h* = 0.0). For all simulations, we assumed a mutation rate of 1.5 × 10^−8^ per site per generation as in Ségurel, Wyman, and Przeworski (2014) and uniform recombination rate of 1 × 10^−8^ crossover events per site per generation (Kong et al. 2010).

We assumed a constant effective population size (*Ne*) of 10,000 for all simulations. We ran burn-ins for 10\**Ne* generations to attain equilibrium prior to outputting the SFS. To explore the impact of simulation variance, we ran 20 total replicates (each resulting in an SFS for 30Mb of coding sequence) and conducted independent demographic and DFE inference with *∂*a*∂*i (Gutenkunst et al. 2009) and Fit*∂*a*∂*i (Kim, Huber, and Lohmueller 2017) on resulting SFS. A schematic figure of our simulation and inference pipeline is shown in **Figure S1**.

### DFE inference

We used the software *∂*a*∂*i (Gutenkunst et al. 2009) and Fit*∂*a*∂*i (Kim, Huber, and Lohmueller 2017) to infer the distributions of selection coefficients of new mutations. Fit*∂*a*∂*i and *∂*a*∂*i rely on the Poisson Random Field model of the SFS (Sawyer and Hartl 1992; Sethupathy and Hannenhalli 2008). Because both demography and selection can change allele frequencies, but we are only interested in quantifying the effects of selection, we accounted for demography by first inferring the demographic parameters (see **Table S2**) using the putatively neutral synonymous sites (the synonymous SFS) with *∂*a*∂*i. We kept the demographic parameters that maximized the likelihood estimates given the data after 30 iterations. Conditioning on resulting demographic parameters, we then inferred selection parameters (DFE) with the nonsynonymous SFS using Fit*∂*a*∂*i. We also collected only MLEs of selection after 30 iterations. Detailed information about the methodology used to infer the DFE and demography is provided in Kim, Huber, and Lohmueller (2017) and Gutenkunst et al. (2009).

#### Fitting the gamma distribution

To assess Fit*∂*a*∂*i’s performance on simulated data, we first inferred the DFE under a gamma distribution. We then calculated the proportion of mutations in different categories of selection coefficient (*s*) by discretizing the gamma distribution into five bins based on the strength of selection coefficients as: neutral: |*s*| = [0, 10^−5^), nearly neutral |*s*| = [10^−5^, 10^−4^), slightly deleterious |*s*| = [10^−4^, 10^−3^), moderately deleterious |*s*| = [10^−3^, 10^−2^), and strongly deleterious |*s*| = [10^−2^, 1].

#### Fitting mixture distributions

We also inferred and evaluated the fit of the DFE on simulated data under distributions of selection coefficients other than the gamma. Specifically, we modeled the DFE using a mixture of distributions as described in Kim, Huber, and Lohmueller (2017): (i) Neu+Gamma: a gamma distribution with a point mass at neutral mutations (|*s*| = [0, 10^−4^]), (ii) Gamma+Let a gamma distribution with a point mass at lethals (|*s*| = [10^−2^, 1]); (iii) Neutral+Gamma+Let, a gamma distribution with a point mass in neutral ([|*s*| = [0, 10^−4^]) and in lethals (|*s*| = [10^−2^, 1]). The different models were compared using Akaike’s Information Criterion (AIC) (Akaike 1992) and results are shown in **Table 1** and **Table 2** under the recessive and additive lethals scenarios, respectively.

#### Profile log-likelihood of the proportion of lethal mutations

To infer the proportion of lethal mutations using Fit*∂*a*∂*i, we performed a log-likelihood profile under the Gamma+Let model. To this end, we fixed the proportion of lethal (*p*_*let*_) at a given value (ranging from 0-0.5) and allowed Fit*∂*a*∂*i to infer the other parameters of the DFE (*α, β*) under the Gamma+Let distribution. For each simulated dataset, we found the value of *p*_*let*_ that gave the maximum likelihood estimate (MLE). We then computed the mean MLE across 20 independent replicates We expect that the closer the mean MLE is to the simulated data’s *p*_*let*_ the greater is the method’s predictive power to accurately infer the true proportion of lethals.

#### Estimation of proportion of lethals through mutation-selection-drift balance

We used mutation-selection-drift balance models coupled with estimates of segregating recessive lethals to estimate the fraction of new mutations that are recessive lethal in humans and in *Drosophila melanogaster*, following the approach from (Amorim et al. 2017). Specifically, this approach leverages the result from (Nei 1968) relating the frequency of a recessive lethal mutation (*q*) to mutation rates (*μ*) and effective population size (*Ne*):

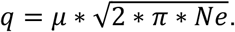

This formula yields the allele frequency at a single locus, whereas estimates of segregating recessive lethals for humans and *D. melanogaster* are genome-wide. Thus, we multiply the resulting allele frequency (*q*) by the length of the coding sequence for each species (*L*) as well as the ratio of nonsynonymous to synonymous sequence length (NS:SYN ratio) (**Table 3**). In doing so, we are therefore assuming that lethal mutations only exist at nonsynonymous sites and no linkage across sites.

To apply this approach for humans, we assumed a mutation rate of 1.5*e* − 8 per site per generation (Ségurel, Wyman, and Przeworski 2014), coding sequence length of 30 Mb (Keightley 2012), and NS:SYN ratio of 2.31:1 (Huber et al. 2017) (see **Table 3** for all parameters). As this model assumes equilibrium, and the human population size has experienced substantial exponential growth (Gravel et al. 2011; Tennessen et al. 2012; Duchen et al. 2013)), we assumed a range of effective population sizes of *Ne* = {20,000, 30,000, 60,000}. These population sizes were selected based on simulations under a human demographic model suggesting that recessive lethal allele frequencies in modern human populations are approximated by an equilibrium effective population size on the order of 20,000-30,000 (see **SI**). However, to account for any potential underestimation of recent exponential growth, we also project model results assuming an equilibrium effective population size of 60,000. We compare predictions from all effective population sizes to a range of segregating recessive lethals of 0.6-1.6 lethals per diploid (Gao et al. 2015; Narasimhan et al. 2016). Although Narasimhan et al. (2016) estimated lethal loss-of-function equivalents––and not recessive lethals––we use their estimate as an upper bound, given that the 0.6 estimate from Gao et al. 2015 is likely a slight underestimate because embryonic lethals were not considered in their inference (Gao et al. 2015).

For *D. melanogaster*, we assumed a mutation rate of 3*e* − 9 per site per generation (Sharp and Li 1989; Keightley et al. 2014), coding sequence length of 22 Mb (Kim et al. 2021), and NS:SYN ratio of 2.85:1 (Huber et al. 2017); see **Table 3** for all parameters). We explored model predictions for several effective population sizes including *Ne* = {5*e*5, 1*e*6, 5*e*6}, a range that encompasses existing estimates (H. Li and Stephan 2006; Duchen et al. 2013; Sheehan and Song 2016; Huber et al. 2017). We compared predictions from this model to an experimentally-estimated range of segregating recessive lethals of 1-3 per diploid (Simmons and Crow 1977; Lynch, Walsh, and Others 1998; McCune et al. 2002).

## Acknowledgements

We thank Bernard Kim and C. Eduardo Guerra Amorim for their helpful discussion on this manuscript. We thank Bernard Kim and Jonathan Mah for their help with the DFE inference using Fit*∂*a*∂*i. This work was supported by NIH grant R35GM119856. EW was a Bruins in Genomics (BIG) student at UCLA supported by NIH grant R25NS115554 to Eleazar Eskin and Roel Ophoff.

## Author Contributions

**Conceptualization:** C.C.K, K.E.L; **Methodology:** E.E.W., C.C.K., M.I.A.C., K.E.L.; **Formal Analysis:** E.E.W., C.C.K., M.I.A.C.; **Resources:** K.E.L.; **Data curation:** E.E.W., C.C.K., M.I.A.C.; **Writing - Original Draft:** C.C.K., M.I.A.C., K.E.L.; **Writing - Review and Editing:** E.E.W., C.C.K., M.I.A.C., K.E.L.; **Visualization:** E.E.W., C.C.K., M.I.A.C., K.E.L. **Supervision:** K.E.L.; **Project administration:** K.E.L.; **Funding acquisition:** K.E.L.

## Data availability

Scripts for simulations and Fit*∂*a*∂*i analysis are available at: https://github.com/emmaewade/Lethals_Project. Scripts for mutation-selection-drift balance results are available at: https://github.com/ckyriazis/lethals_scripts.

## References

Agarwal, Ipsita, and Molly Przeworski. 2021. “Mutation Saturation for Fitness Effects at Human CpG Sites.” eLife 10 (November). https://doi.org/10.7554/eLife.71513.

Agrawal, Aneil F., and Michael C. Whitlock. 2011. “Inferences About the Distribution of Dominance Drawn From Yeast Gene Knockout Data.” Genetics 187 (2): 553–66.

Amorim, Carlos Eduardo G., Ziyue Gao, Zachary Baker, José Francisco Diesel, Yuval B. Simons, Imran S. Haque, Joseph Pickrell, and Molly Przeworski. 2017. “The Population Genetics of Human Disease: The Case of Recessive, Lethal Mutations.” PLoS Genetics 13 (9): e1006915.

Ballinger, Mallory A., and Mohamed A. F. Noor. 2018. “Are Lethal Alleles Too Abundant in Humans?” Trends in Genetics: TIG.

Bank, Claudia, Gregory B. Ewing, Anna Ferrer-Admettla, Matthieu Foll, and Jeffrey D. Jensen. 2014. “Thinking Too Positive? Revisiting Current Methods of Population Genetic Selection Inference.” Trends in Genetics: TIG 30 (12): 540–46.

Bittles, A. H., and J. V. Neel. 1994. “The Costs of Human Inbreeding and Their Implications for Variations at the DNA Level.” Nature Genetics 8 (2): 117–21.

Boyko, Adam R., Scott H. Williamson, Amit R. Indap, Jeremiah D. Degenhardt, Ryan D. Hernandez, Kirk E. Lohmueller, Mark D. Adams, et al. 2008. “Assessing the Evolutionary Impact of Amino Acid Mutations in the Human Genome.” PLoS Genetics 4 (5): e1000083.

Campos José L., Daniel L. Halligan, Penelope R. Haddrill, and Brian Charlesworth. 2014. “The Relation between Recombination Rate and Patterns of Molecular Evolution and Variation in Drosophila Melanogaster.” Molecular Biology and Evolution 31 (4): 1010–28.

Castellano, David, Marta Coronado-Zamora, Jose L. Campos, Antonio Barbadilla, and Adam Eyre-Walker. 2015. “Adaptive Evolution Is Substantially Impeded by Hill–Robertson Interference in Drosophila.” Molecular Biology and Evolution 33 (2): 442–55.

Castellano, David, Moisès Coll Macià, Paula Tataru, Thomas Bataillon, and Kasper Munch. 2019. “Comparison of the Full Distribution of Fitness Effects of New Amino Acid Mutations Across Great Apes.” Genetics 213 (3): 953–66.

Cavassim, Maria Izabel A., Stig U. Andersen, Thomas Bataillon, and Mikkel Heide Schierup. 2021. “Recombination Facilitates Adaptive Evolution in Rhizobial Soil Bacteria.” Molecular Biology and Evolution 38 (12): 5480–90.

Dobzhansky, T., B. Spassky, and N. Spassky. 1954. “Rates of Spontaneous Mutation in the Second Chromosomes of the Sibling Species, Drosophila Pseudoobscura and Drosophila Persimilis.” Genetics 39 (6): 899–907.

Dubinin, N. P. 1946. “On Lethal Mutations in Natural Populations.” Genetics 31 (January): 21–38.

Duchen, Pablo, Daniel Zivkovic, Stephan Hutter, Wolfgang Stephan, and Stefan Laurent. 2013. “Demographic Inference Reveals African and European Admixture in the North American Drosophila Melanogaster Population.” Genetics 193 (1): 291–301.

Dukler, Noah, Mehreen R. Mughal, Ritika Ramani, Yi-Fei Huang, and Adam Siepel. 2021. “Extreme Purifying Selection against Point Mutations in the Human Genome.” bioRxiv. https://doi.org/10.1101/2021.08.23.457339.

Elyashiv, Eyal, Kevin Bullaughey, Shmuel Sattath, Yosef Rinott, Molly Przeworski, and Guy Sella. 2010. “Shifts in the Intensity of Purifying Selection: An Analysis of Genome-Wide Polymorphism Data from Two Closely Related Yeast Species.” Genome Research 20 (11): 1558–73.

Eyre-Walker, Adam, and Peter D. Keightley. 2009. “Estimating the Rate of Adaptive Molecular Evolution in the Presence of Slightly Deleterious Mutations and Population Size Change.” Molecular Biology and Evolution 26 (9): 2097–2108.

Eyre-Walker, Adam, Megan Woolfit, and Ted Phelps. 2006. “The Distribution of Fitness Effects of New Deleterious Amino Acid Mutations in Humans.” Genetics 173 (2): 891–900.

Fuller, Zachary L., Jeremy J. Berg, Hakhamanesh Mostafavi, Guy Sella, and Molly Przeworski. 2019. “Measuring Intolerance to Mutation in Human Genetics.” Nature Genetics 51 (5): 772–76.

Galtier, Nicolas. 2016. “Adaptive Protein Evolution in Animals and the Effective Population Size Hypothesis.” PLoS Genetics 12 (1): e1005774.

Galtier, Nicolas, and Marjolaine Rousselle. 2020. “How Much Does Ne Vary Among Species?” Genetics 216 (2): 559–72.

Gao, Ziyue, Darrel Waggoner, Matthew Stephens, Carole Ober, and Molly Przeworski. 2015. “An Estimate of the Average Number of Recessive Lethal Mutations Carried by Humans.” Genetics 199 (4): 1243–54.

Gilbert, Kimberly J., Stefan Zdraljevic, Daniel E. Cook, Asher D. Cutter, Erik C. Andersen, and Charles F. Baer. 2022. “The Distribution of Mutational Effects on Fitness in Caenorhabditis Elegans Inferred from Standing Genetic Variation.” Genetics 220 (1). https://doi.org/10.1093/genetics/iyab166.

Gravel, Simon, Brenna M. Henn, Ryan N. Gutenkunst, Amit R. Indap, Gabor T. Marth, Andrew G. Clark, Fuli Yu, Richard A. Gibbs, 1000 Genomes Project, and Carlos D. Bustamante. 2011. “Demographic History and Rare Allele Sharing among Human Populations.” Proceedings of the National Academy of Sciences of the United States of America 108 (29): 11983–88.

Gutenkunst, Ryan N., Ryan D. Hernandez, Scott H. Williamson, and Carlos D. Bustamante. 2009. “Inferring the Joint Demographic History of Multiple Populations from Multidimensional SNP Frequency Data.” PLoS Genetics 5 (10): e1000695.

Haller, Benjamin C., and Philipp W. Messer. 2019. “SLiM 3: Forward Genetic Simulations Beyond the Wright–Fisher Model.” Molecular Biology and Evolution 36 (3): 632–37.

Halligan, Daniel L., and Peter D. Keightley. 2003. “How Many Lethal Alleles?” Trends in Genetics: TIG 19 (2): 57–59.

Huang, Xin, Alyssa Lyn Fortier, Alec J. Coffman, Travis J. Struck, Megan N. Irby, Jennifer E. James, José E. León-Burguete, Aaron P. Ragsdale, and Ryan N. Gutenkunst. 2021. “Inferring Genome-Wide Correlations of Mutation Fitness Effects between Populations.” Molecular Biology and Evolution 38 (10): 4588–4602.

Huber, Christian D., Arun Durvasula, Angela M. Hancock, and Kirk E. Lohmueller. 2018. “Gene Expression Drives the Evolution of Dominance.” Nature Communications 9 (1): 2750.

Huber, Christian D., Bernard Y. Kim, Clare D. Marsden, and Kirk E. Lohmueller. 2017. “Determining the Factors Driving Selective Effects of New Nonsynonymous Mutations.” Proceedings of the National Academy of Sciences of the United States of America 114 (17): 4465–70.

Hvilsom, Christina, Yu Qian, Thomas Bataillon, Yingrui Li, Thomas Mailund, Bettina Sallé, Frands Carlsen, et al. 2012. “Extensive X-Linked Adaptive Evolution in Central Chimpanzees.” Proceedings of the National Academy of Sciences of the United States of America 109 (6): 2054–59.

Jouganous, Julien, Will Long, Aaron P. Ragsdale, and Simon Gravel. 2017. “Inferring the Joint Demographic History of Multiple Populations: Beyond the Diffusion Approximation.” Genetics 206 (3): 1549–67.

Kardos, Marty, Ellie E. Armstrong, Sarah W. Fitzpatrick, Samantha Hauser, Philip W. Hedrick, Joshua M. Miller, David A. Tallmon, and W. Chris Funk. 2021. “The Crucial Role of Genome-Wide Genetic Variation in Conservation.” Proceedings of the National Academy of Sciences of the United States of America 118 (48). https://doi.org/10.1073/pnas.2104642118.

Keightley, Peter D. 2012. “Rates and Fitness Consequences of New Mutations in Humans.” Genetics 190 (2): 295–304.

Keightley, Peter D., and Adam Eyre-Walker. 2007. “Joint Inference of the Distribution of Fitness Effects of Deleterious Mutations and Population Demography Based on Nucleotide Polymorphism Frequencies.” Genetics 177 (4): 2251–61.

Keightley, Peter D., Rob W. Ness, Daniel L. Halligan, and Penelope R. Haddrill. 2014. “Estimation of the Spontaneous Mutation Rate per Nucleotide Site in a Drosophila Melanogaster Full-Sib Family.” Genetics 196 (1): 313–20.

Kim, Bernard Y., Christian D. Huber, and Kirk E. Lohmueller. 2017. “Inference of the Distribution of Selection Coefficients for New Nonsynonymous Mutations Using Large Samples.” Genetics 206 (1): 345–61.

Kim, Bernard Y., Jeremy R. Wang, Danny E. Miller, Olga Barmina, Emily Delaney, Ammon Thompson, Aaron A. Comeault, et al. 2021. “Highly Contiguous Assemblies of 101 Drosophilid Genomes.” eLife 10 (July). https://doi.org/10.7554/eLife.66405.

Kong, Augustine, Gudmar Thorleifsson, Daniel F. Gudbjartsson, Gisli Masson, Asgeir Sigurdsson, Aslaug Jonasdottir, G. Bragi Walters, et al. 2010. “Fine-Scale Recombination Rate Differences between Sexes, Populations and Individuals.” Nature 467 (7319): 1099–1103.

Koufopanou, Vassiliki, Susan Lomas, Isheng J. Tsai, and Austin Burt. 2015. “Estimating the Fitness Effects of New Mutations in the Wild Yeast Saccharomyces Paradoxus.” Genome Biology and Evolution 7 (7): 1887–95.

Li, Haipeng, and Wolfgang Stephan. 2006. “Inferring the Demographic History and Rate of Adaptive Substitution in Drosophila.” PLoS Genetics 2 (10): e166.

Li, Yingrui, Nicolas Vinckenbosch, Geng Tian, Emilia Huerta-Sanchez, Tao Jiang, Hui Jiang, Anders Albrechtsen, et al. 2010. “Resequencing of 200 Human Exomes Identifies an Excess of Low-Frequency Non-Synonymous Coding Variants.” Nature Genetics 42 (11): 969–72.

Lynch, Michael, Bruce Walsh, and Others. 1998. “Genetics and Analysis of Quantitative Traits.” http://www.invemar.org.co/redcostera1/invemar/docs/RinconLiterario/2011/febrero/AG_8.pdf.

Ma, Xin, Joanna L. Kelley, Kirsten Eilertson, Shaila Musharoff, Jeremiah D. Degenhardt, André L. Martins, Tomas Vinar, et al. 2013. “Population Genomic Analysis Reveals a Rich Speciation and Demographic History of Orang-Utans (Pongo Pygmaeus and Pongo Abelii).” PloS One 8 (10): e77175.

McCune, Amy R., Rebecca C. Fuller, Allisan A. Aquilina, Robert M. Dawley, James M. Fadool, David Houle, Joseph Travis, and Alexey S. Kondrashov. 2002. “A Low Genomic Number of Recessive Lethals in Natural Populations of Bluefin Killifish and Zebrafish.” Science 296 (5577): 2398–2401.

Morton, N. E., J. F. Crow, and H. J. Muller. 1956. “AN ESTIMATE OF THE MUTATIONAL DAMAGE IN MAN FROM DATA ON CONSANGUINEOUS MARRIAGES.” Proceedings of the National Academy of Sciences of the United States of America 42 (11): 855–63.

Moutinho, Ana Filipa, Fernanda Fontes Trancoso, and Julien Yann Dutheil. 2019. “The Impact of Protein Architecture on Adaptive Evolution.” Molecular Biology and Evolution 36 (9): 2013–28.

Narasimhan, Vagheesh M., Karen A. Hunt, Dan Mason, Christopher L. Baker, Konrad J. Karczewski, Michael R. Barnes, Anthony H. Barnett, et al. 2016. “Health and Population Effects of Rare Gene Knockouts in Adult Humans with Related Parents.” Science 352 (6284): 474–77.

Nei, M. 1968. “The Frequency Distribution of Lethal Chromosomes in Finite Populations.” Proceedings of the National Academy of Sciences of the United States of America 60 (2): 517–24.

Pérez-Pereira, Noelia, Armando Caballero, and Aurora García-Dorado. 2022. “Reviewing the Consequences of Genetic Purging on the Success of Rescue Programs.” Conservation Genetics 23 (1): 1–17.

Ragsdale, Aaron P. 2021. “Can We Distinguish Modes of Selective Interactions Using Linkage Disequilibrium?” bioRxiv. https://doi.org/10.1101/2021.03.25.437004.

Sawyer, S. A., and D. L. Hartl. 1992. “Population Genetics of Polymorphism and Divergence.” Genetics. https://doi.org/10.1093/genetics/132.4.1161.

Ségurel, Laure, Minyoung J. Wyman, and Molly Przeworski. 2014. “Determinants of Mutation Rate Variation in the Human Germline.” Annual Review of Genomics and Human Genetics 15 (June): 47–70.

Sethupathy, Praveen, and Sridhar Hannenhalli. 2008. “A Tutorial of the Poisson Random Field Model in Population Genetics.” Advances in Bioinformatics. https://doi.org/10.1155/2008/257864.

Sharp, P. M., and W. H. Li. 1989. “On the Rate of DNA Sequence Evolution in Drosophila.” Journal of Molecular Evolution 28 (5): 398–402.

Sheehan, Sara, and Yun S. Song. 2016. “Deep Learning for Population Genetic Inference.” PLoS Computational Biology 12 (3): e1004845.

Simmons, M. J., and J. F. Crow. 1977. “Mutations Affecting Fitness in Drosophila Populations.” Annual Review of Genetics 11: 49–78.

Tennessen, Jacob A., Abigail W. Bigham, Timothy D. O’Connor, Wenqing Fu, Eimear E. Kenny, Simon Gravel, Sean McGee, et al. 2012. “Evolution and Functional Impact of Rare Coding Variation from Deep Sequencing of Human Exomes.” Science 337 (6090): 64–69.

