## Supplementary Information for "Quantifying the fraction of new mutations that are recessive lethal"

**Comparing predictions from mutation-selection-drift balance to simulations**

As noted in the main text, mutation-selection-drift balance models assume that populations are in demographic equilibrium. As this assumption does not hold for humans or *D. melanogaster*, we ran forward-in-time simulations using SLiM 3 (Haller and Messer 2019) under a non-equilibrium demographic model to assess the impact of violations of this model assumption. Due to the large effective population size estimates for *D. melanogaster* (H. Li and Stephan 2006; Duchen et al. 2013; Sheehan and Song 2016; Huber et al. 2017), which are computationally infeasible for forward-in-time simulations, we restricted this analysis to humans.

For our simulations, we assumed a human demographic model inferred by Jouganous et al. (2017), which is an update of the Gravel et al. (2011) model. This model consists of an ancestral population size in the African population of $Ne = 11{,}273$ followed by growth to $Ne = 23{,}721$, with European and Asian populations diverging and undergoing a bottleneck followed by exponential growth (see simulation scripts for all model parameters). We ran burn-ins at $Ne = 11{,}273$ for $5000$ generations, which was sufficient for recessive lethal mutations to reach equilibrium (**Fig. S6**). We assumed identical genomic parameters in these simulations as those used for our mutation-selection-drift balance analysis (**Table 3**) and assumed a recessive lethal fraction of 0.5%. To visualize results, we plotted simulation results against model predictions under mutation-selection-drift balance at 20 generation intervals.

Here, we find that recessive lethal mutations increase rapidly to their equilibrium frequency, reaching equilibrium within roughly ~500 generations of a population size change (**Fig. S6**). At the onset of the out-of-Africa bottleneck, recessive lethal alleles are rapidly purged in European and Asian populations observed at a frequency consistent with their much smaller effective population sizes. During subsequent exponential growth in these populations, recessive lethal alleles rapidly increase in frequency, but at a rate that does not quite track the rapid pace of population growth (**Fig. S6**). At the conclusion of exponential growth, there are ~1.15 recessive lethal alleles per diploid in the European population and ~1.35 recessive lethal alleles in the Asian population. These values are consistent with an equilibrium effective population size in the European population of ~21,000 and in the Asian population of ~29,000, despite the actual present-day effective sizes numbering 39,530 and 83,289, respectively (**Fig. S6**).

Overall, we conclude that recessive lethal allele frequencies in modern-day human populations can be approximated using an equilibrium effective population size on the order of 20,000-30,000. More broadly, these results demonstrate that non-equilibrium demography does greatly impact recessive lethal allele frequencies. However, these effects appear to be muted in humans by the counteracting effects of the out-of-Africa bottleneck and subsequent exponential growth. Finally, these results also demonstrate the need to examine a wide range of equilibrium effective population sizes for our mutation-selection-balance results in *D. melanogaster*, given the wide range of estimated effective population sizes in the species and evidence for bottlenecks and exponential growth in many populations (Duchen et al. 2013; Sheehan and Song 2016; Huber et al. 2017; H. Li and Stephan 2006).

**Table S1. The proportion of synonymous, nonsynonymous, and nonsynonymous lethal mutations across simulated datasets.** Within each dataset, deleterious (nonsynonymous) and neutral (synonymous) mutations occurring are modeled at a ratio of 2.31:1. The percentages of each mutation type (synonymous, nonsynonymous, lethal) are described. $p_{let}$ corresponds to the proportion of nonsynonymous mutations that are lethal for each dataset.

| **Dataset ($p_{let}$)** | **% mutations synonymous** | **% mutations nonsynonymous from gamma DFE (69.8 * (1 - $p_{let}$))** | **% nonsynonymous mutations lethal (69.8 * $p_{let}$)** |
|---|---|---|---|
| 0 | 30.2 | 69.8 | 0 |
| 0.01 | 30.2 | 69.102 | 0.698 |
| 0.05 | 30.2 | 66.31 | 3.49 |
| 0.10 | 30.2 | 62.82 | 6.98 |

**Table S2. Demographic and population genetic parameters used in simulations**
Population genetics parameters used to scale the DFE in terms of $s$. Parameters are as follows: $N_{chrm}$ denotes the number of haploid chromosomes; $\theta_S$ is the mutation rate at synonymous sites; $L_{NS}/L_S$ is the ratio of possible nonsynonymous to synonymous sites; $\theta_{NS}$ is the population scaled mutation rate of nonsynonymous sites, $L_S$ is the total count of synonymous sites, $L_{NS}$ is the total count of nonsynonymous sites, $N_{anc}$ denotes the ancestral population size which was inferred as $Nanc = \theta_S/(4 * \mu * L_{NS})$ and the $\boldsymbol{\theta_{NS}}$ was computed as $\theta_{NS} = \theta_S * L_{NS}/L_S$. nu is the ratio of contemporary to ancient population size, and T is the time in the past at which size change occurred (in 2Na units).

| Dataset | $\boldsymbol{N_{chrm}}$ | $\theta_S$ | $\boldsymbol{L_{NS}/L_S}$ | $\theta_{NS}$ | $\mu$ | $\boldsymbol{L_{NS} + L_S}$ | $\boldsymbol{L_S}$ | $\boldsymbol{L_{NS}}$ | Nanc | nu | T |
|---|---|---|---|---|---|---|---|---|---|---|---|
| Simulation | 10; 100; 1000 | 7,941 | 2.31 | 18,344 | 1.5e-8 | 0.44 MB | 0.13 MB | 0.31 MB | 10,000 | 1 | 0 |

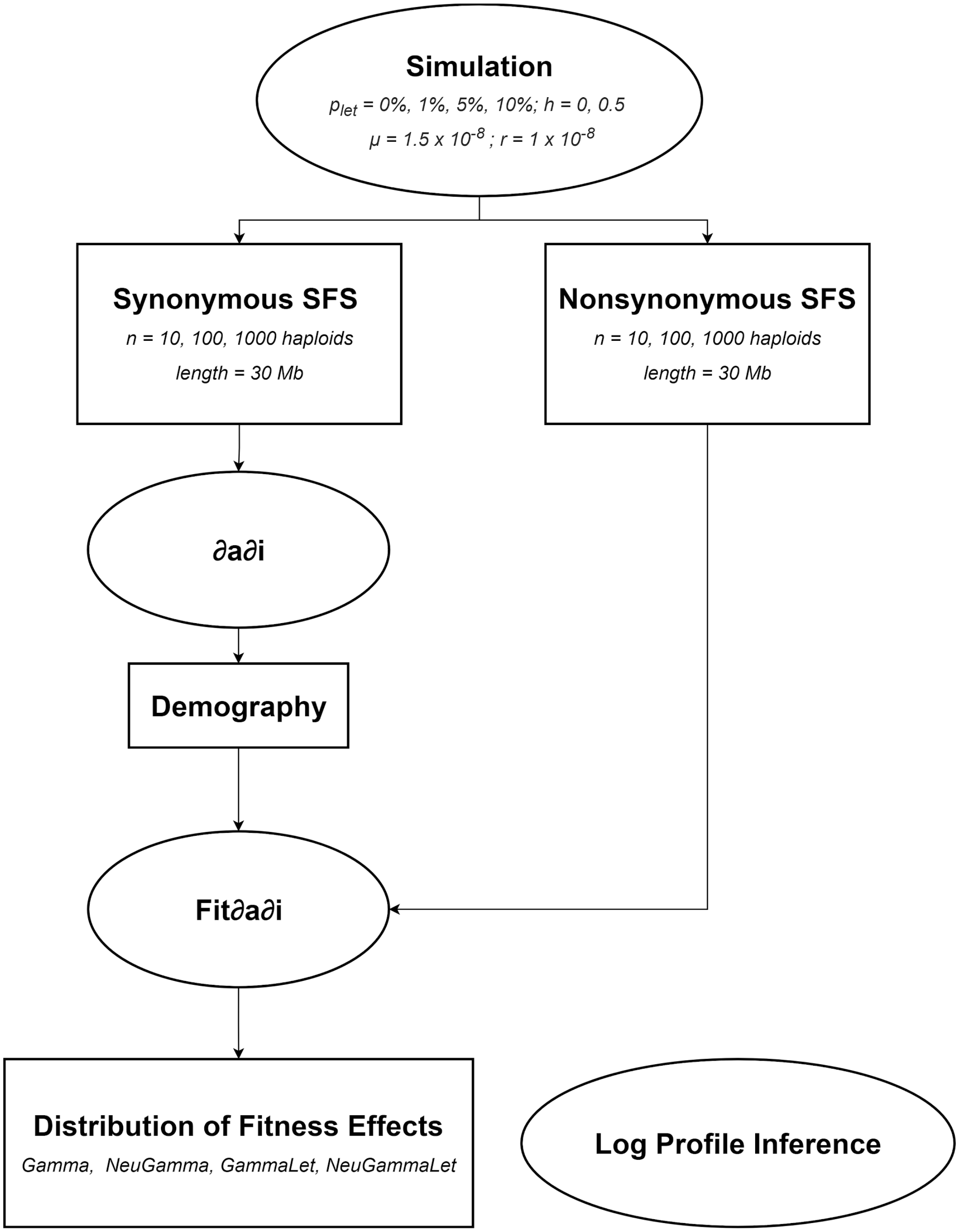


**Figure S1. Schematic representation of all simulation frameworks used in the present study**. Forward-in-time simulations were run using the software SLiM 3 (Haller and Messer 2019). Simulations were conducted with increments of proportions of lethals ($p_{let}$) and increments of sample size (number of haploids). From simulations, the synonymous (neutral mutations) and nonsynonymous (lethal + deleterious mutations) site frequency spectra (SFS) were computed and used for demographic and DFE inference, respectively. To approximate the parameters observed in humans, a mutation rate ($\mu$) of $1.5 \times 10^{-5}$ and a constant recombination rate of $1.0 \times 10^{-8}$ were assumed as previously described (Ségurel, Wyman, and Przeworski 2014; Kong et al. 2010).

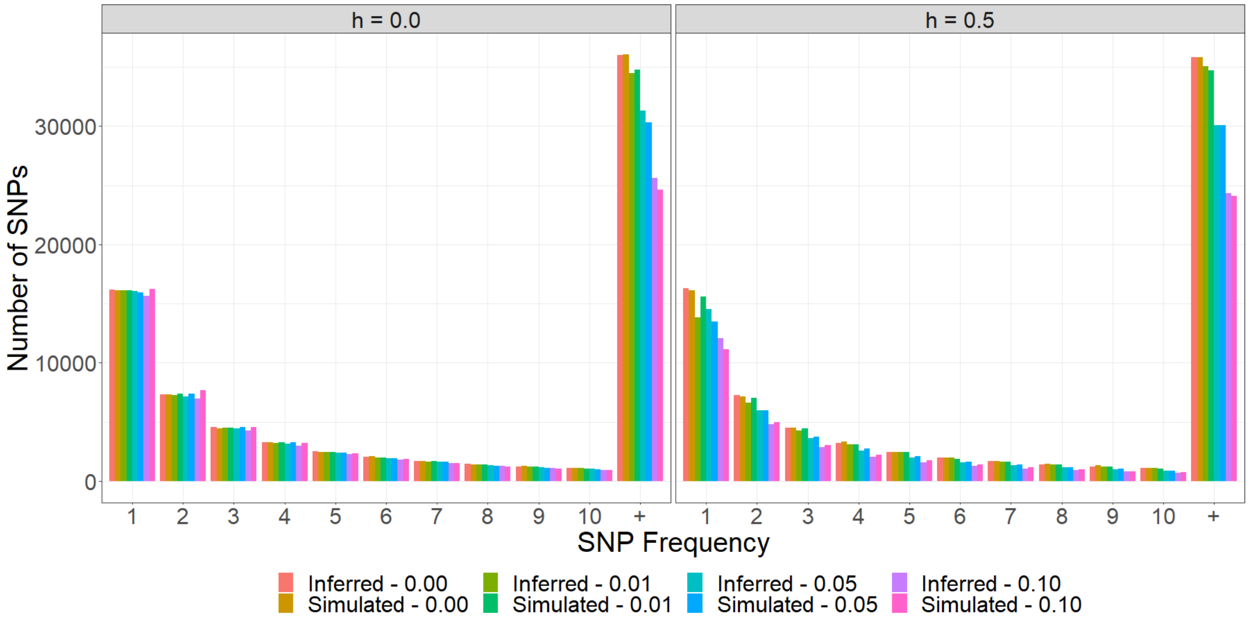


**Figure S2. Simulated and inferred nonsynonymous SFS.** Comparison between the nonsynonymous SFS from the simulated datasets (simulated) and the SFS inferred from the best-fitting DFE (inferred) under the gamma DFE model and inclusion of recessive lethal (left panel, $h$=0) and additive lethal (right panel, $h$=0.5) mutations. Colors represent different datasets based on the proportion of lethals: salmon and dark yellow correspond to the inferred and simulated SFS under 0% of lethals; shades of green under 1% of lethals, shades of blue under 5% lethals, and pink and purple under 10% of lethals.

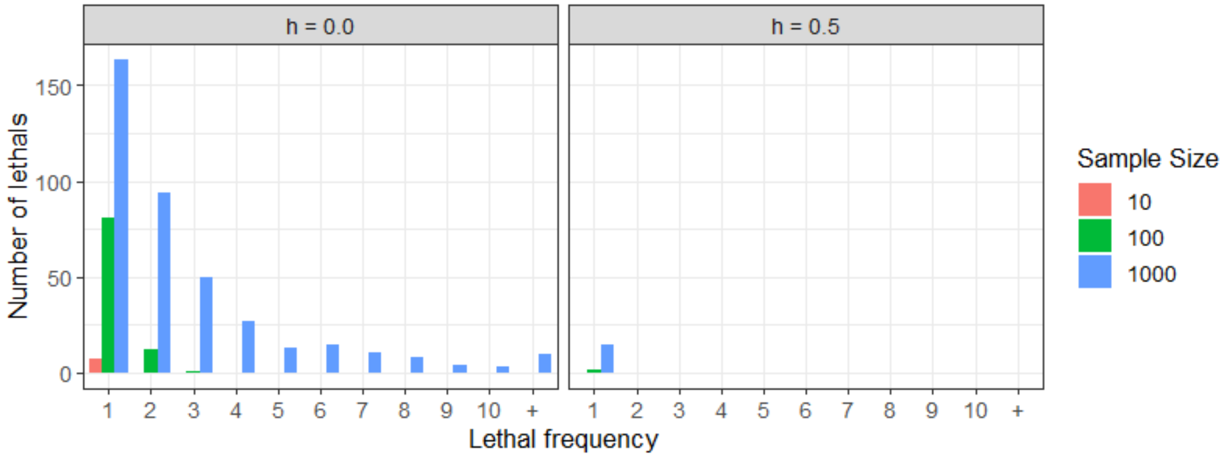


**Figure S3. The SFS when 1% of nonsynonymous mutations are lethal for different sample sizes assuming different dominance coefficients.** Left panel corresponds to the SFS of lethals assuming that lethal mutations are recessive, while the right panel corresponds to the SFS of additive lethal mutations.

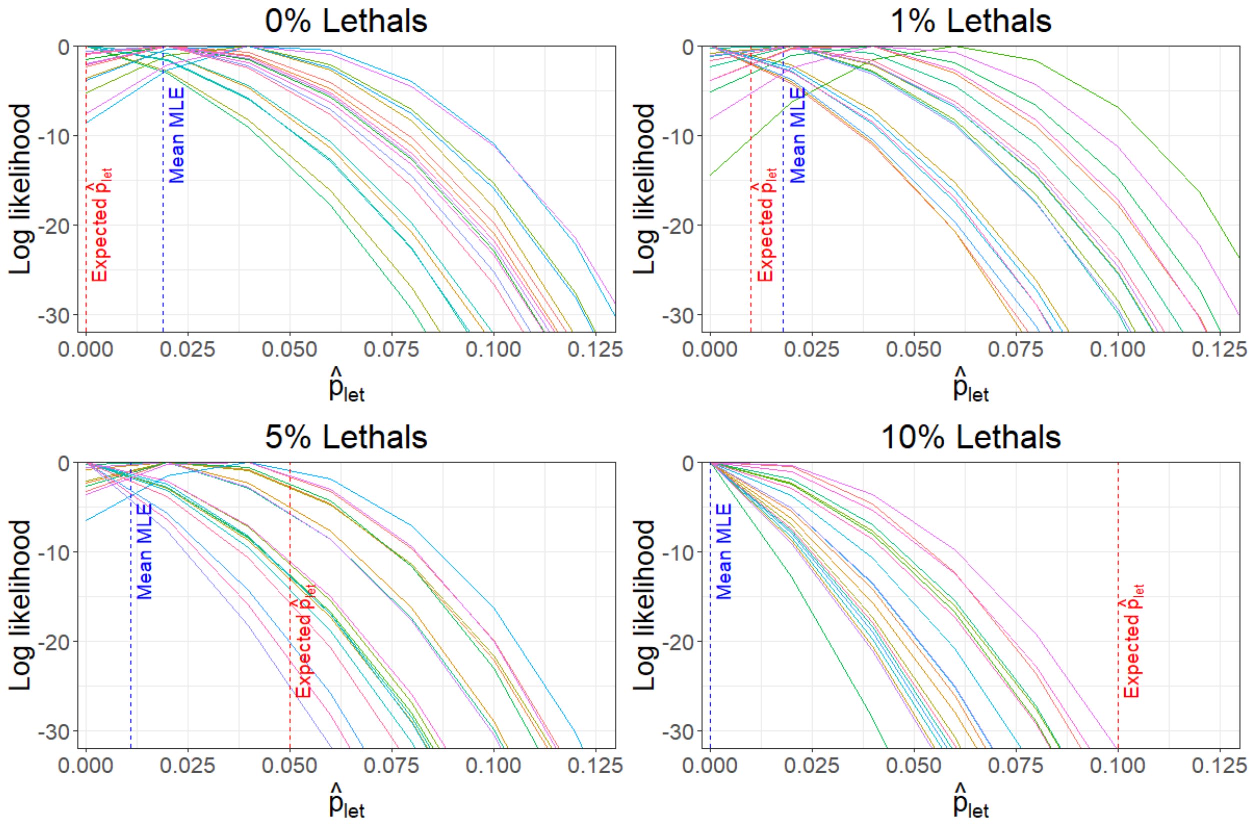


**Figure S4. Evaluating the performance of Fit∂a∂i under a gamma + recessive lethal DFE.** The x-axis indicates the proportion of recessive lethals, and the y-axis denotes the log-likelihood for each value of $p_{let}$. Each line corresponds to a simulation replicate under different increments of recessive lethals. Red dashed lines correspond to the expected (true) lethal proportion under each lethal simulated class, while blue dashed lines represent the mean maximum likelihood estimate across 20 independent replicates for each lethal class.

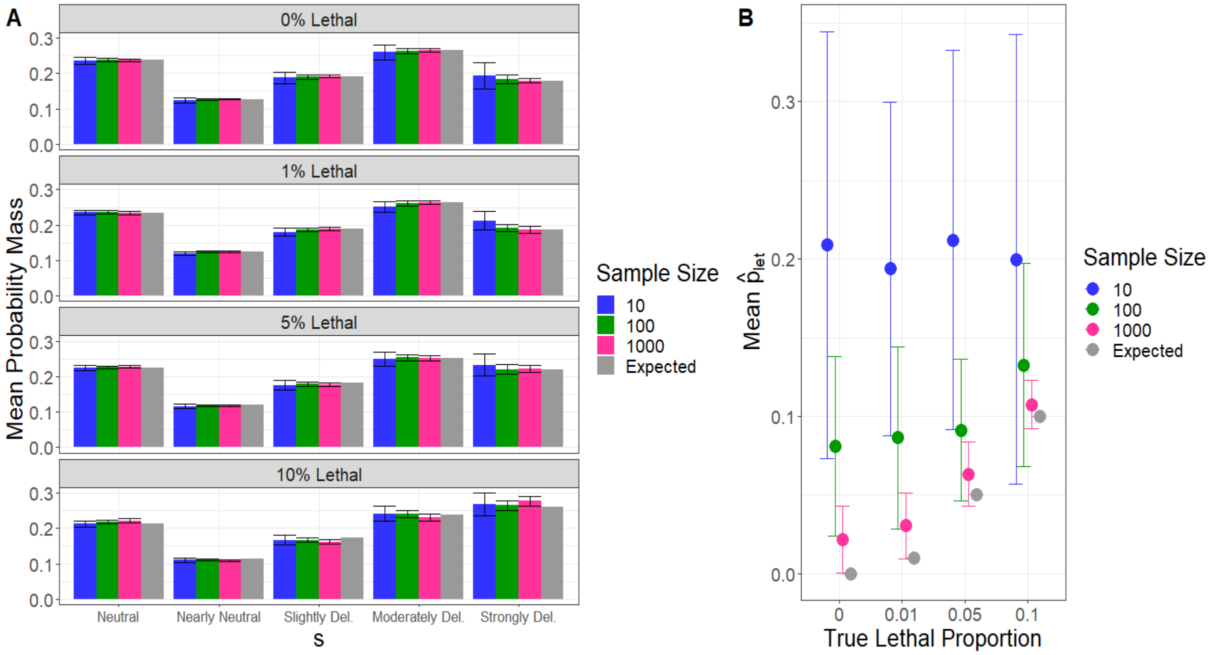


**Figure S5. Inferred DFE and proportion of additive lethals ($\hat{p}_{let}$) depending on the sample size and on the portion of additive lethals mutations simulated. A.** The mean probability mass obtained with the gamma parameters for each DFE category. The DFE categories are defined as neutral ($|s| = [0;\ 10^{-5})$), nearly neutral ($|s| = [10^{-5};\ 10^{-4})$), slightly deleterious ($|s| = [10^{-4};\ 10^{-3})$), moderately deleterious ($|s| = [10^{-3};\ 10^{-2})$), and strongly deleterious ($|s| = [10^{-2}, 1]$). Error bars correspond to the range of inferred proportions obtained across 20 simulation replicates. **B.** The mean proportion of lethals inferred ($\hat{p}_{let}$) with the Gamma+Let model and different sample sizes. The gray dots correspond to the expected proportion of lethals ($p_{let}$). Error bars correspond to the range of values of $\hat{p}_{let}$ inferred from 20 simulation replicates.

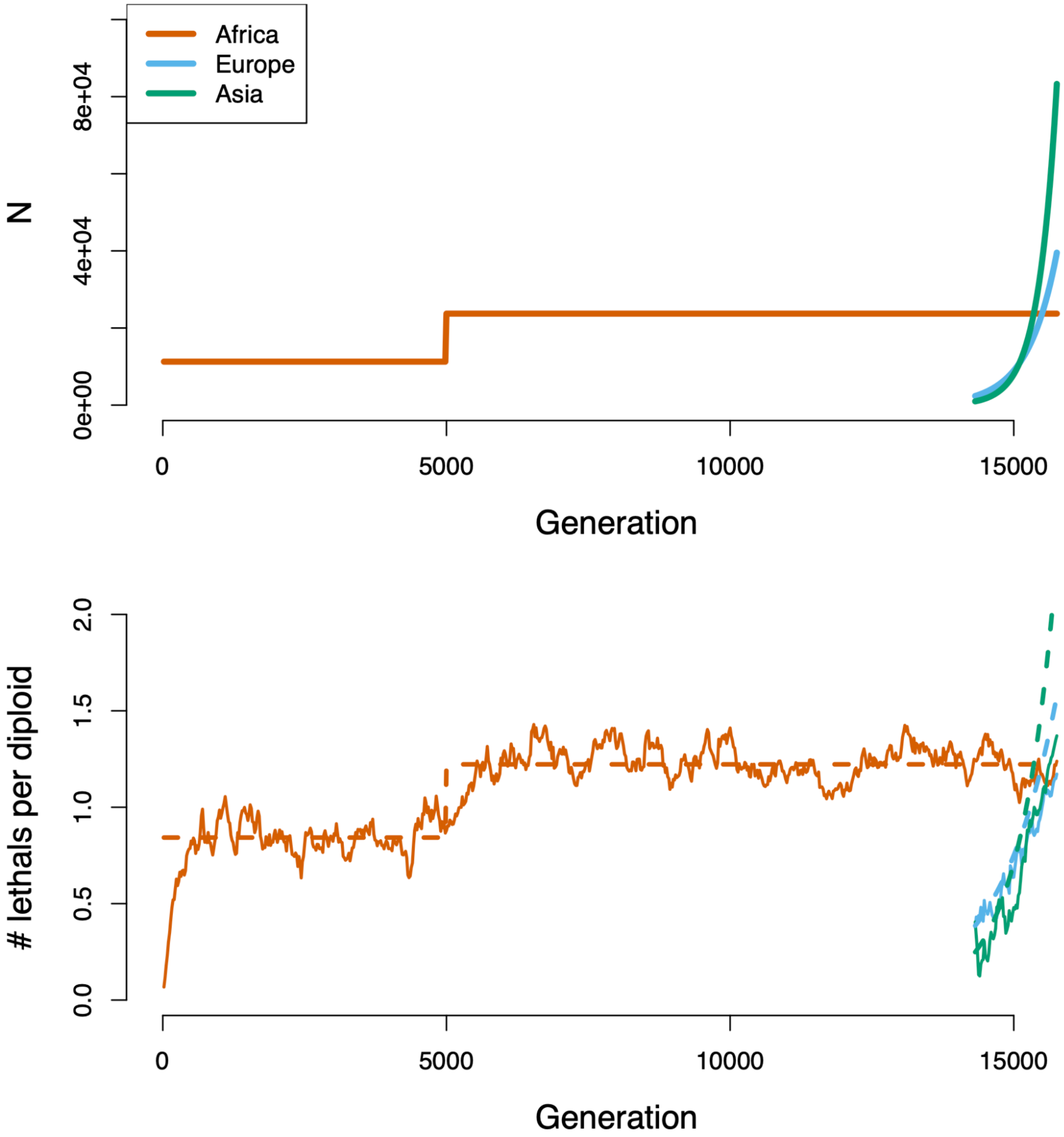


**Figure S6: Comparison of analytical and simulation predictions of lethals under a human demographic model from Jouganous 2017**. Top panel depicts simulated effective population sizes for the African, European, and Asian populations. Bottom panel depicts resulting numbers of recessive lethals per diploid in solid line and analytical prediction from mutation-selection-drift balance in dashed line. Note that the simulated lethals closely track the analytical prediction, with the exception of during rapid exponential growth in the European and Asian populations.
